# Logic models to predict continuous outputs based on binary inputs with an application to personalized cancer therapy

**DOI:** 10.1101/036970

**Authors:** Theo Knijnenburg, Gunnar Klau, Francesco Iorio, Mathew Garnett, Ultan McDermott, Ilya Shmulevich, Lodewyk Wessels

## Abstract

Mining large datasets using machine learning approaches often leads to models that are hard to interpret and not amenable to the generation of hypotheses that can be experimentally tested. Finding ‘actionable knowledge’ is becoming more important, but also more challenging as datasets grow in size and complexity. We present ‘Logic Optimization for Binary Input to Continuous Output’ (LOBICO), a computational approach that infers small and easily interpretable logic models of binary input features that explain a binarized continuous output variable. Although the continuous output variable is binarized prior to optimization, the continuous information is retained to find the optimal logic model. Applying LOBICO to a large cancer cell line panel, we find that logic combinations of multiple mutations are more predictive of drug response than single gene predictors. Importantly, we show that the use of the continuous information leads to robust and more accurate logic models. LOBICO is formulated as an integer programming problem, which enables rapid computation on large datasets. Moreover, LOBICO implements the ability to uncover logic models around predefined operating points in terms of sensitivity and specificity. As such, it represents an important step towards practical application of interpretable logic models.

## Introduction

Regression and classification models are important tools for researchers in various fields. The application of these many-to-one mapping models is two-fold. First, they can be used for prediction. The output value or class of a (new) case can be predicted by applying the inferred mapping to the input variables of the case. Second, they inform us about the relationship between the input and the output. They specify how the input variables are (mathematically) interacting with each other to produce the output variable. The usefulness of the second application is, however, limited by the power of the human intellect. We suggest that the interpretation of these many-to-one mapping models is of utmost, yet undervalued, importance in many research fields.

This also holds for computational biology, where often, a multitude of molecular and genomic data is used to explain or predict a biological or clinical phenotype. Single predictor models are generally not accurate enough, reflecting the importance of acknowledging the interaction between biological components. On the other hand, machine learning approaches, such as Elastic Net [1] and Random Forests [2] produce complex multi-predictor models that are hard to interpret and not amenable to the generation of hypotheses that can be experimentally tested. As a consequence, such models are not likely to further our understanding of biology. There is an urgent need for approaches that build small, interpretable, yet accurate models that capture the interplay between biological components and explain the phenotype of interest.

In this study, we have developed such a modeling framework to explain drug response of cancer cell lines using gene mutation data. Our approach, ‘Logic Optimization for Binary Input to Continuous Output’ (LOBICO) infers small and easily interpretable logic models of gene mutations (binary input variables) that explain the observed sensitivity to anticancer drugs in the cell lines (continuous output).

The contributions of our approach are three-fold: First, the continuous information of the output variable is retained in the logic mapping. The output variable is binarized, which facilitates its interpretation, yet the distances of the continuous values to the binarization threshold are used in the inference. Second, LOBICO provides the user with the option to include constraints on the model performance that allows the identification of logic models around operating points predefined in terms of sensitivity and specificity. This enables tailoring of the model to, for example, clinical applications where the severity of diseases or side effects of the treatment dictate a desired level of specificity or sensitivity. Third, the logic mapping is formulated as an integer linear programming problem (ILP). This means that advanced ILP solvers can be used to find an optimal logic mapping fast enough to apply LOBICO to large and complex datasets without the need to tune parameters.

Our work is similar in spirit to logic regression (LR) [3, 4], sparse combinatorial inference (SCI) [5], Markov logic networks [6, 7], combinatorial association logic (CAL) [8] and genetic programming for association studies (GPAS) [9], which all employ combinatorial logic to explicitly incorporate interactions in their models. LOBICO differentiates itself from these approaches by its direct emphasis on interpretability. This is in contrast with the linearly weighted sums of logic functions as inferred by LR or the posterior probabilities of predictors in the model averaged across an ensemble of many solutions as inferred by SCI. Moreover, LOBICO includes constraints on statistical performance criteria, such as a minimum specificity, which is a novel feature not available in any of the other approaches.

Here, we demonstrate LOBICO by application to a large cancer cell line panel, where the goal is to explain drug response based on binary mutation data of a set of genes [10]. We investigate whether logic models perform better than single-gene predictors, and put genes that co-occur in logic models in the context of known cancer pathways. We assess whether using continuous output values provides benefit in terms of robustness and performance above the use of binarized data, which is usually the starting point for logical analysis of data [11]. Finally, we explore the power and flexibility of the ILP formulation by inferring logic models around different operating points in the receiver-operator-characteristic (ROC) space. We show that the ability to find robust and interpretable models with a predefined balance between false positives and false negatives is an important step towards practical application of these models.

## Results

### Sensitivity to anticancer drugs as a logic combination of gene mutations

We used LOBICO to find the logic combinations of mutations that best explain the response of cancer cell lines to anticancer drugs. This analysis was based on a panel of 714 cancer cell lines from more than 50 tumor types for which the binary mutation status of 54 known cancer genes was obtained [10]. A gene was called mutated when it had a point mutation, a small insertion or deletion as determined by capillary sequencing, or when it was highly amplified or homozygously deleted based on copy number arrays. Additionally, we included the presence or absence of 6 known oncogenic gene fusions across these cell lines, resulting in a binary data matrix with a total of 60 features. The cell lines in the panel were screened against 142 anticancer drugs. The half-maximal inhibitory concentrations at 72 hours (IC50s) obtained in this screen were used to represent the drug response and served as the continuous output variables. LOBICO was applied to the IC50s of each drug separately.

Figure 1 provides a schematic overview of the application of LOBICO to this dataset using the IC50s of the EGFR/ERBB2 inhibitor Afatinib as the output variable. For Afatinib, IC50s were obtained for 642 of the 714 cell lines. Although LOBICO uses the continuous values, it is also necessary to define a binarization threshold for the output variable. This threshold is used to divide the cell lines into two classes; the sensitive cell lines and the resistant cell lines. LOBICO finds the optimal logic function of binary mutation features that minimizes the error, which is defined as the sum of the weighted misclassified cell lines. A misclassification occurs when a sensitive cell line is predicted by the model to be resistant or vice versa. The weight is proportional to the distance to the binarization threshold. Consequently, misclassification of cell lines close to the binarization threshold does not considerably affect the optimization criterion, whereas there is a large penalty for misclassifying cell lines that are extremely sensitive or resistant to the drug. By default, the total weight associated with samples of each class is normalized in order to balance class importance. This is especially important for unbalanced classes as is often the case for the cell line panel, where we observed that, for most drugs, the bulk of cell lines are not affected, and only a small percentage (5-15% typically) end up in the class of sensitive cell lines. The optimization procedure is formulated as an integer linear programming (ILP) problem, which provides an efficient way to find optimal solutions. See **Methods Section** for more details.

**Figure 1.**
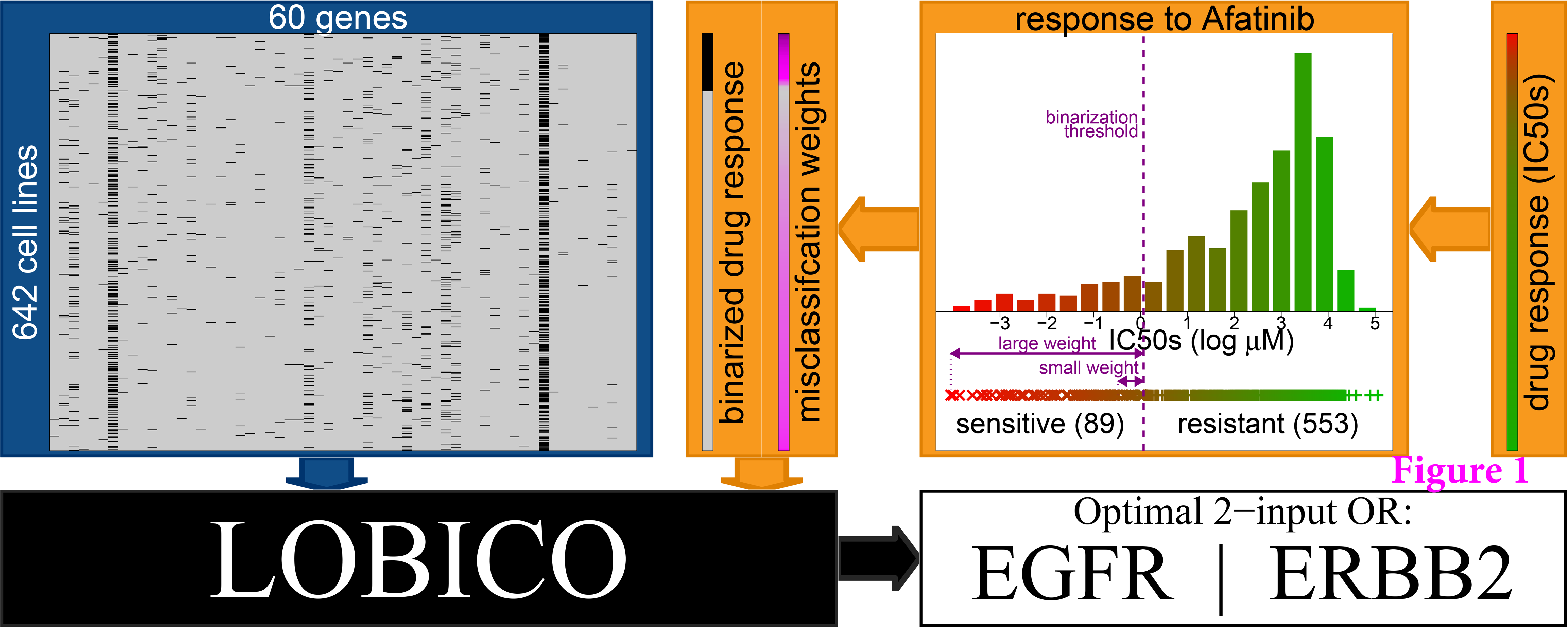
| Workflow of LOBICO. LOBICO has two main inputs: (1) a binary matrix of samples by features (depicted in the blue box). Here, the binary matrix contains the mutation status of 60 cancer genes measured across 642 cancer cell lines. (2) a continuous vector with a value for each of the samples (depicted in the orange boxes). In this case, the vector contains the IC50 of each cell line in response to Afatinib, an EGFR/ERRB2 inhibitor. The continuous vector is transformed into a binary vector and a sample-specific weight vector using a binarization scheme. Particularly, the IC50s are binarized using a threshold leading to a set of sensitive and a set of resistant cell lines. The distances of the original IC50s to the binarization threshold are represented in the weight vector, which is normalized per class. Then, LOBICO finds the optimal logic model of features (gene mutations) that minimizes the total weight of misclassified samples (cell lines). In this case, the optimal 2-input OR logic formula is ‘EGFR OR ERBB2’ (depicted in the white box).

### Drug response is explained by multiple mutations

LOBICO finds logic models that are described in the disjunctive normal form (DNF), a standard notation in which every logic function can be expressed. The DNF is parameterized by two parameters: *K*, the number of disjuncts, and *M*, the number of terms per disjunct. We applied LOBICO with eight different parameter settings, i.e. from simple single-predictor models (*K*=1, *M*=1) to more complex multi-predictor models (*K*>1, *M*>1). See Table 1 for examples of the eight models. A stratified 10-fold cross-validation (CV) strategy was employed to select the appropriate model complexity, i.e. the one with the lowest CV error.

**Table 1.**
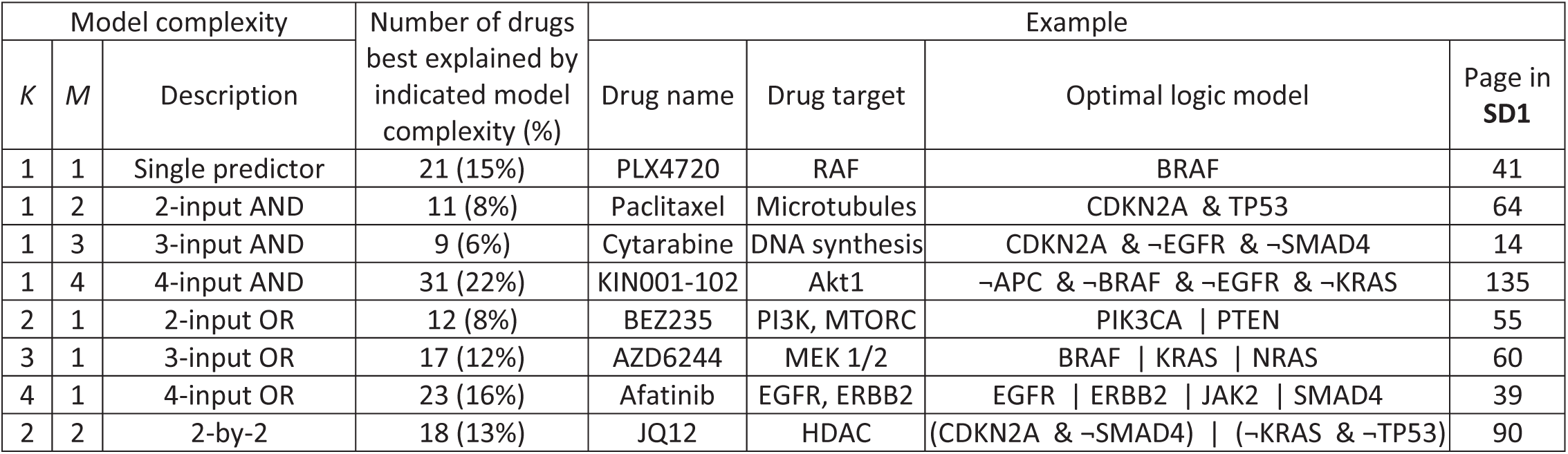
Table 1 | Overview of the optimal logic model complexity as determined by crossvalidation across the set of 142 drugs. For each of the 8 model complexities (columns 1-3), this table states the number of drugs best explained by the indicated model complexity (column 4), as well an example of one drug, including drug name (column 5), drug target (column 6), the optimal logic formula (column 7) and the page in Supplementary Data 1 with the standard LOBICO visualization of the inferred logic models for the indicated drug (column 8).

Application across all 142 drugs revealed that the large majority of drugs (>85%) is best explained by multi-predictor models (Table 1, Supplementary Data 1 and 2). In many cases the multi-predictor model selected by CV was substantially better than the single-predictor model (Figure 2, Supplementary Figure 1). For example, LOBICO inferred a 4-input OR model for the MEK1/2 inhibitor PD-0325901, which had a CV error of 0.21 while the best single input model only reached 0.34. The 4-input OR model included the genes BRAF, NRAS, KRAS and HRAS. Thus, cell lines with a BRAF mutation or a mutation in NRAS, KRAS or HRAS (members of the RAS protein family) are predicted to be sensitive to this drug. Since BRAF and RAS are directly upstream of MEK in the MEK-ERK pathway, this association is easily understood from pathway biology. In general, OR models will increase the number of cell lines (or patient population) that are predicted to respond to a drug. On the other hand, AND models refine the predictions. For example, LOBICO inferred that cell lines with a mutation in both tumor suppressors TP53 and CDKN2A specifically respond to the microtubule inhibitor Paclitaxel (Taxol). These results strengthen the notion that gene mutations should not be considered in isolation, but in relationship to one another, if they are to be used as prognostic biomarkers or predictors of therapy response.

**Figure 2.**
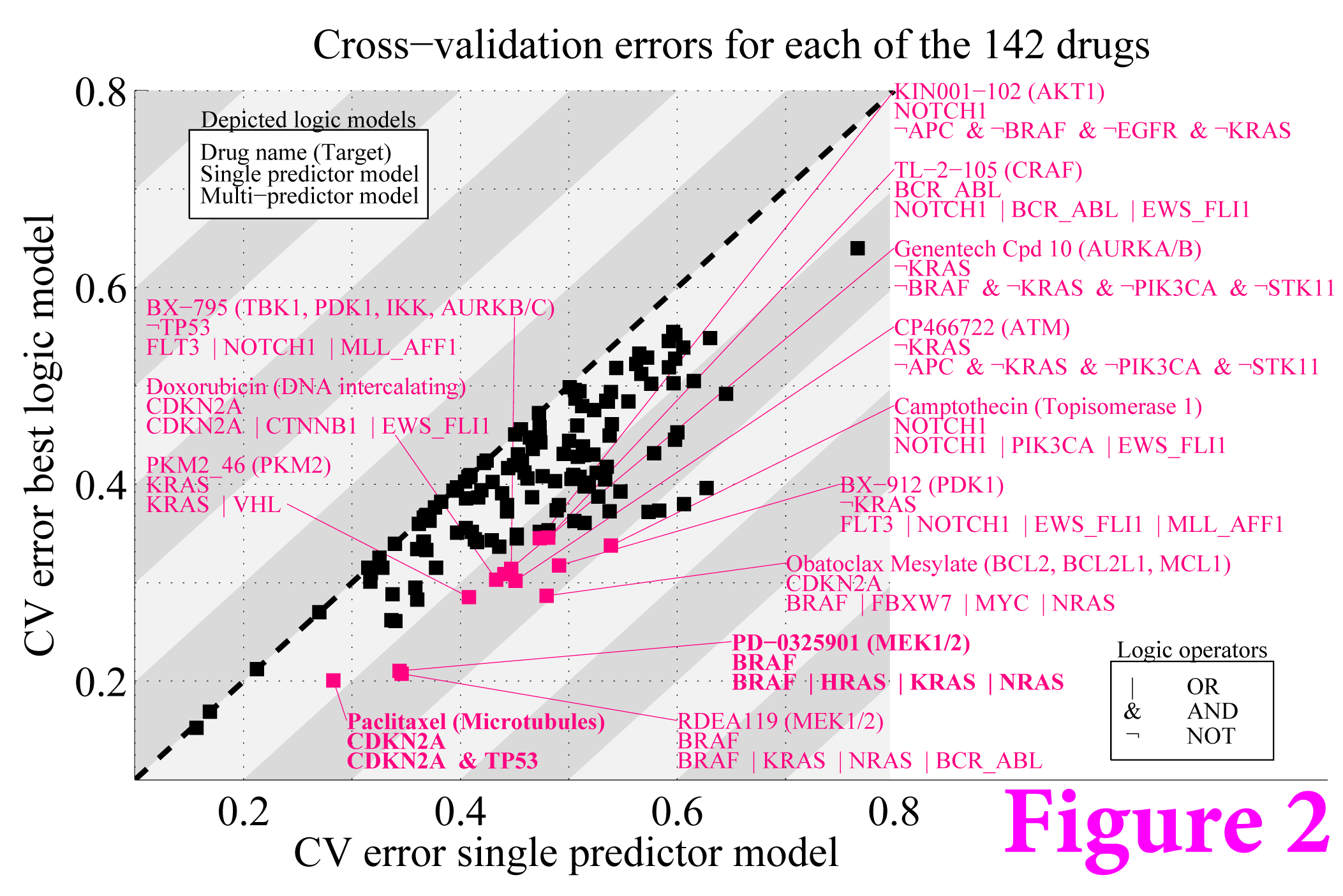
| Multi-predictor models outperform single-predictor models. Scatter plot with the 10-fold cross-validation (CV) error for single predictor models (x-axis) and the best (lowest CV error) multi-predictor model (y-axis). Each point represents one of the 142 drugs. Multi-predictor models that have a CV error lower than 0.35 and at least a 25% improvement upon the single predictor model are highlighted in magenta. The two examples discussed in the text are highlighted in bold typeface.

### Use of continuous output yields more robust and accurate models

After LOBICO was successfully employed to infer sensible models of drug response, we set out to quantify the contribution of the use of the sample-specific weights. In other words, is it advantageous to use the distances from the continuous IC50s to the binarization threshold as sample-specific misclassification penalties? Or, would simply using binarized data as is done in standard logical data analysis [11] as well as in a recent application to cancer cell line drug screening data [12] be equally informative?

To this end, we analyzed logic models under slightly different binarization thresholds. Specifically, we shifted the default threshold (*t*=0.05) slightly downwards (*t*=0.03) and upwards (*t*=0.07) leading to, on average, 24% less and 22% more sensitive cell lines, respectively. See **Methods Section** for details. Then, we ran LOBICO across these binarization thresholds with and without the sample-specific weights. The logic models for a drug were inferred using the model complexity (defined by *K* and *M*) selected by CV for that drug in the standard setting, i.e. with the sample-specific weights and *t*=0.05. We conjectured that small changes in the binarization thresholds should only moderately affect the inferred logic models, i.e. the models should be robust against these small changes.

We evaluated this notion by using the correlation of feature importance (FI) scores across the binarization thresholds as a measure of robustness. Thus, the degree of change in the FI scores is inversely proportional to the robustness of the model. The FI score for a feature is defined as the increase in error when the feature is left out of the inferred logic model (Equation 9). The FI scores are not only based on the optimal logic model, but also on other good (suboptimal) logic models that are identified by the ILP solver. (See **Methods Section** for details.) Figure 3a depicts the FI scores of the 60 gene mutation features for the PI3K/mTOR inhibitor BEZ235. From this panel it is clear that two features, PTEN and PI3KCA, are important in explaining BEZ235 response. More importantly, when sample-specific weights are employed, the FI remains robust across different values of the discretization threshold.

**Figure 3.**
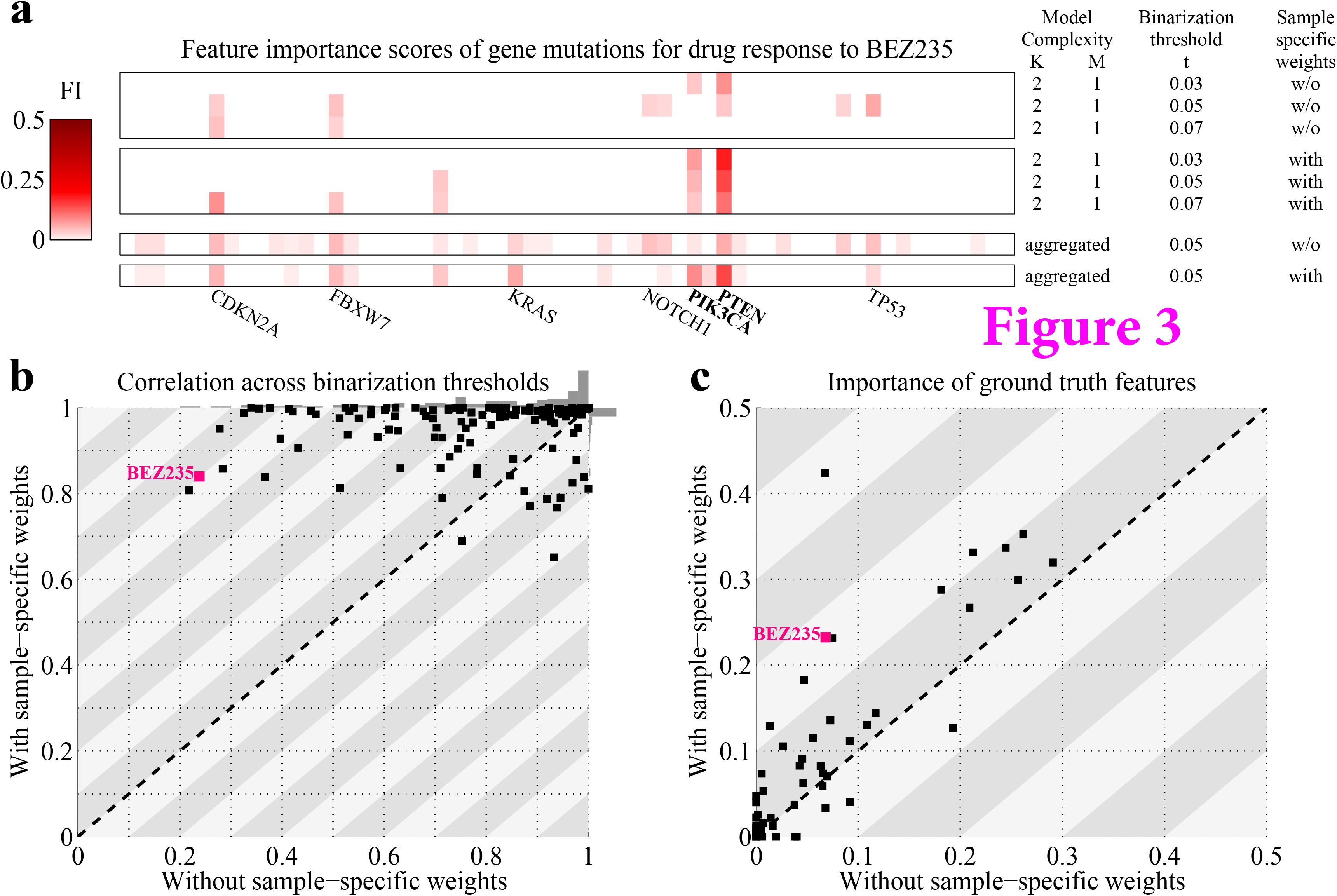
| LOBICO’s use of continuous output leads to robust and accurate models. **a)** Heatmaps depleting the feature Importance (FI) scores across the 60 gene mutations for the logic models inferred to explain the drug response to the PI3K/mTOR inhibitor BEZ235. The upper heatmap represents FI scores for the 2-input OR model (*K*=2, *M*=1) using three different binarization thresholds for logic models with binarized output, i.e. not using the sample-specific weights. The middle of the three heatmaps represents the same FI scores, but for logic models with continuous output, i.e. using the sample-specific weights. The bottom two heatmaps depict FI scores aggregated across all model complexities, using the standard binarization threshold (*t*=0.05), for both the logic models with and without the sample-specific weights. The labels of the gene mutations with a large FI in any of these heatmaps are printed below. The ‘ground truth’ features, i.e. the expected or annotated targets of this drug, PTEN and PIK3CA, are printed in bold. **b)** Scatter plot with the average Pearson correlation coefficients of the similarity of FI scores across the binarization thresholds for inferred logic models without (x-axis) and with (y-axis) the sample-specific weights. Each point represents one of the 142 drugs. The correlation scores are computed using the model-complexity-specific FI scores. The grey bars on top and to the right of the scatter plot represent histograms of these correlation scores for models without and with the sample-specific weights, respectively. c) Scatter plot with the importance of the ground truth features for inferred logic models without (x-axis) and with (y-axis) the sample-specific weights. Each point represents one of the 49 drugs, for which ground truth features were available. The importance scores of the ground truth features were derived from aggregated FI scores.

Also across the remaining drugs the use of the continuous output via sample-specific weights resulted in a substantially smaller variation in the FI scores across the binarization thresholds (Figure 3b). Particularly, the FI scores for the logic models across the three thresholds had an average Pearson correlation coefficient larger than 0.75 for all but two drugs (99% of the drugs), and 106 drugs (75% of the drugs) had a correlation larger than 0.95. This was in stark contrast to the logic models based without the sample-specific weights, where 93 drugs (65%) had a correlation larger than 0.75 and only 44 (31%) had a correlation larger than 0.95. From this observation we conclude that the use of the sample-specific weights based on the continuous output makes the inferred logic models less sensitive to small changes in the dataset, i.e. more robust.

We also assessed the robustness of the logic models across the CV training folds. Here, we observed a similar pattern, although the difference between using and not using sample-specific weights was less pronounced (**Supplementary Note 1** and Supplementary Figure 2). Logic models inferred from the randomly sampled subsets of cell lines, i.e. the CV folds, showed more variability in feature importance scores than the models inferred with slight changes in the binarization threshold. This is not surprising, since the in- or exclusion of samples, especially those far away from the binarization threshold, can have a large effect on the optimization function (Equation 1) and thus the inferred optimal logic model. One such example in our dataset is represented by the drug Bicalutamide, which has the poorest CV performance (point in the top right in Figure 2). Here, the cell line HT-3 has an extremely low IC50, much lower than all other sensitive cell lines (see **Supplementary Data 1**). This results in a disproportionally large weight for this sample (its weight is 0.21), whereas the total weight of all 657 cell lines, for which IC50s have been obtained, is 1. This analysis hints at an important weakness of our approach: Outliers, i.e. samples with erroneously large or small output values, have a large effect on the inferred logic model. In general, it is thus very important that outliers are detected and removed before performing a LOBICO analysis.

Next, we employed a ‘ground truth’ mapping between drugs and mutations to gauge whether the use of continuous output variables led to more accurate models. This mapping, which is based on the annotation of the drug targets and expert knowledge, links 49 drugs to one or more of the 60 gene mutation features (**Supplementary Data 2**). For each of these 49 drugs, we analyzed the FI scores of the ground truth features. In this case, we used ‘aggregated’ FI scores, which were not just based on the optimal and suboptimal models from the model complexity with the lowest CV error, but also on the other model complexities that have a CV error equal or smaller than the CV error for the single-predictor model (*K*=1, *M*=1). These aggregated FI scores provide a more comprehensive landscape of the importance of gene mutations in explaining drug response (Figure 3a).

For 19 of the 49 drugs, the total aggregated FI scores of the ground truth features were not larger than 0.05, both for the logic models with as well as without the sample-specific weights (Figure 3c). In these cases, the ground truth features did not explain the observed variation in drug response. For 26 of the remaining 30 drugs, the logic models with sample specific weights had larger FI scores for the ground truth features versus only 4 for the logic models without sample specific weights. Moreover, for 7 drugs, the use of the sample specific weights increased the importance of the ground truth features by at least 0.1. These 7 drugs included the PI3K/mTOR inhibitor BEZ235, 2 BRAF inhibitors and 4 small molecule inhibitors that target PDGFRA and KIT mutants, two of which also target FLT3 mutants, according to the ground truth mapping. Amongst the sensitive cell lines for these drugs, those with the smallest IC50s, i.e. the most sensitive ones, indeed harbored mutations in the ground truth features. This explains the increased accuracy of the logic models with sample-specific weights of retrieving the ground truth features.

The use of sample-specific weights is a prominent feature of LOBICO that sets it apart from traditional methods that use binarized data. The results presented in this section demonstrate that the use of the continuous output leads to more robust and accurate logic models.

### LOBICO finds logic models around a user-defined operating point

A binary classifier can make two kinds of mistakes: false positives (Type I errors) and false negatives (Type II errors). The trade-off between the false positive rate (FPR, 1- specificity) and false negative rate (FNR, 1-sensitivity) defines the operating point of the classifier. The practical application of a binary classifier in any domain is determined by its operating point, the prevalence of positives in the population of interest and the costs, financial or otherwise, that are associated with false positives, false negatives and the test itself.

For example, many medical screening tests, which are relatively inexpensive and generally non-invasive, have high sensitivity and are useful for ‘ruling out’ patients that test negative. For example, a mammogram to screen for breast cancer has a sensitivity of 80% and a specificity of 90% [13]. (It should be noted that periodic screening amounts to a large total number of false positives as evidenced by the fact that half of the women screened in the US will receive a false positive mammogram in any 10-year period [14].)

The implementation of LOBICO as an integer programming problem allows one to add constraints to ensure that solutions meet predefined statistical performance criteria, such as a minimum specificity. We have applied lower bounds on the sensitivity and specificity by adding additional constraints to the ILP (see Equations 10 and 11). If a solution is found, the corresponding logic model is guaranteed to meet these constraints. By setting various lower bounds on the sensitivity and specificity we can probe the receiver-operator-characteristic (ROC) space. In that way we can uncover logic models possibly employing different features at different operating points. Note that this analysis is quite different from the classical ROC analysis, where the same model is evaluated with different thresholds on the output parameter.

For the 25 drugs with the lowest CV error in the original analysis, we ran LOBICO with an array of sensitivity and specificity constraints covering the complete ROC space in intervals of 0.05. The optimal solution for each combination of sensitivity and specificity was again determined by CV. Note that the inclusion of the constraints could lead to different model complexities having the smallest CV error. The Pareto front of solutions is formed by the logic models that perform best in terms of the tradeoff between sensitivity and specificity. This front is equivalent to the ROC curve, although the model at each operating point is different. We observed a large variation in the logic models across this curve, both in terms of the genes that were present in the models as well as the model complexity (**Supplementary Data 3**).

Figure 4a depicts the logic models in the ROC space for a single drug: MEK inhibitor AZD6244 (brand name Selumetinib), which is currently in clinical trials. Mutations in the gene BRAF clearly play an important role in explaining the drug response in the cell line panel, since BRAF is found in almost all solutions in the ROC space. The most specific solution near the bottom-left of the curve (FPR<5%) states that BRAF mutants that neither have a CDKN2A nor a TP53 mutation are sensitive to this drug. This high specificity solution can be termed a ‘rule in’ solution, because due to the low FPR, cell lines (or, potentially in the future, patients) that carry a BRAF mutation but are CDKN2A and TP53 wild-type are very likely to respond to this drug. However, this solution explains only slightly more than 20% of the sensitive cell lines (true positive rate (TPR)≈20%). Around the middle of the ROC curve, we observed a 3-input OR solution consisting of BRAF, KRAS and NRAS, the latter two being part of the RAS protein family, which is directly upstream of BRAF, MEK and ERK in the MAPK pathway. This solution has a TPR slightly below 60% and a FPR<30%. At the top-right of the ROC curve, LOBICO provided a high sensitivity solution (TPR>95%) that did not contain BRAF. This 4-input AND model predicts that cell lines that are wild-type (WT) for PIK3CA, RB1, STK11 mutations and that do not contain the EWS-FLI1 gene fusion, will respond to the drug. Another way to interpret this solution is that cell lines that have a mutation in either PIK3CA, RB1, STK11 or that contain the EWS-FLI1 gene fusion are resistant to the drug with a very high degree of certainty (FPR<5% for identifying resistant cell lines). This high sensitivity solution can be termed a ‘rule out’ solution, as cell lines (or potentially patients) that satisfy this rule will most likely not respond to this drug.

**Figure 4.**
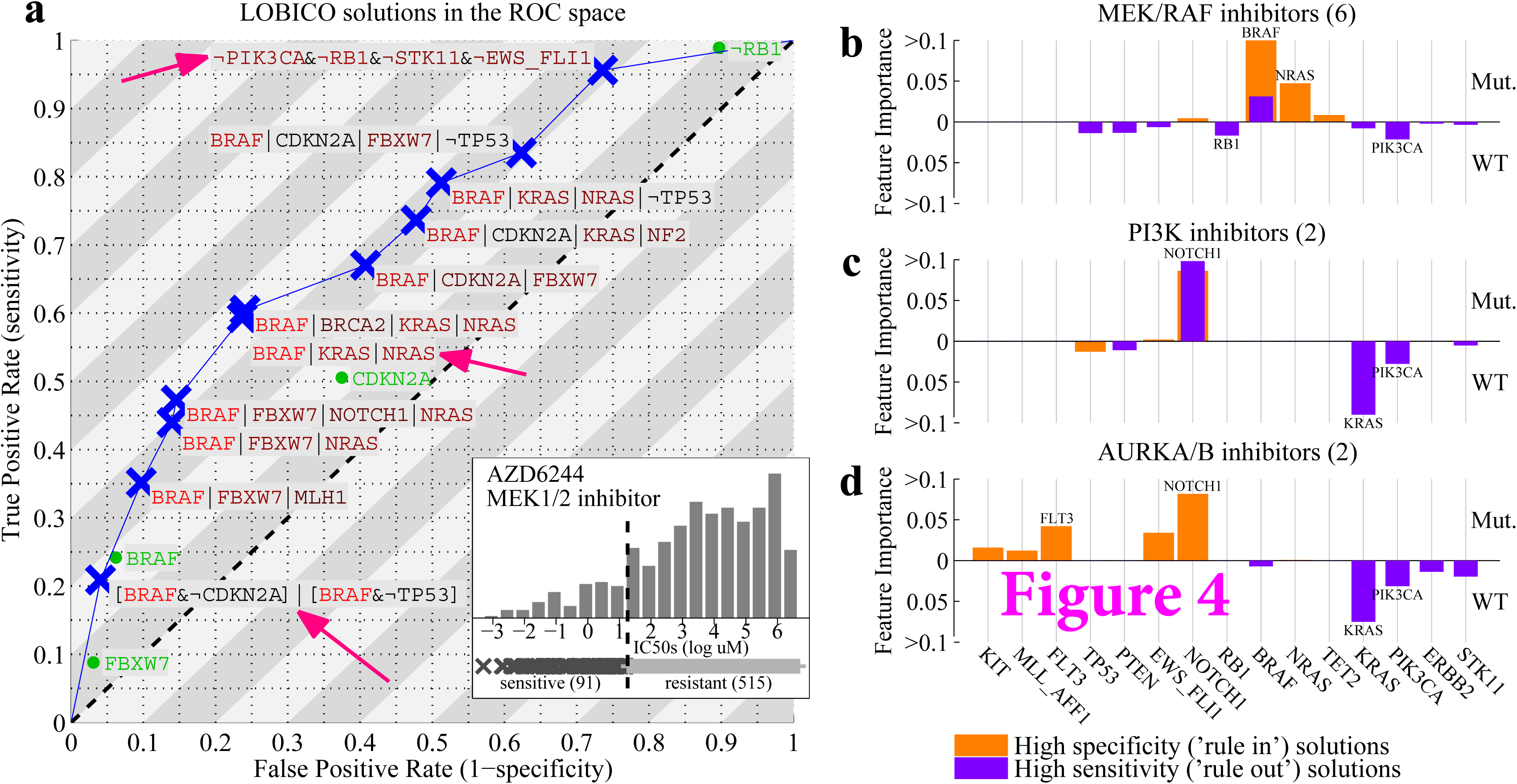
| LOBICO finds solutions at different operating points. **a)** ROC space with LOBICO solutions to explain drug sensitivity to the MEK1/2 inhibitor AZD6244. Blue crosses indicate the TPR and FPR at which the solution was found. The logic formula of the solutions is printed next to the blue crosses. The color of the genes in a formula indicate their FI. Colors range from black (moderately important) to bright red (highly important). For comparison, the best single-predictor solutions are visualized in green. Pink arrows point to solutions discussed in the text. The inlay depicts the histogram of IC50s for AZD6244 together with the binarization threshold, which divides the cell lines into 91 cell lines that are sensitive to AZD6244 and 515 that are resistant. **b)** Average FI scores for a group of 6 MEK/RAF inhibitors (including AZD6244), for high specificity solutions (orange) and high sensitivity solutions (magenta). High specificity solutions were defined as solutions with FPR<10%. Conversely, high sensitivity solutions were defined as solutions with TPR>90%. The FI scores of all solutions on the Pareto front (ROC curve) that met these respective criteria across the six drugs were averaged. We distinguished between positive terms, indicating mutations (Mut.) and negated terms, indicating wild-type (WT). The two genes with the highest average FI score as mutants were printed at the top of their FI bar. The two genes with the highest average FI score as wild-types were printed at the bottom of their FI bar. c), d) Similar to b), but for a group of two PI3K inhibitors and a group of two AURKA/B inhibitors, respectively.

Across the 25 drugs, there were 6 inhibitors of MEK or RAF, 2 inhibitors of Phosphoinositide 3-kinase (PI3K) and 2 inhibitors of Aurora kinase (AURK). The logic models across the ROC space displayed high similarity within these three groups (Supplementary Figure 3). Figure 4b, c and d show the average FI scores of the high-specificity (‘rule in’) solutions and the high-sensitivity (‘rule out’) solutions. As already observed with the MEK-inhibitor AZD6244, mutations in BRAF and NRAS are indicative of drug response with high specificity for the MEK/RAF inhibitors. Conversely, the high importance scores for wild-type RB1 and PIK3CA in the high sensitivity solutions indicate that RB1 and PIK3CA mutants are resistant to these inhibitors. These observations fit within the current ideas on oncogenic signaling, as mutations in the tumor suppressor RB1 and oncogene PIK3CA can lead to uncontrolled cell growth by activating pathways other than the MAPK pathway that these inhibitors are targeting [15, 16]. For the PI3K and AURK inhibitors we observed that cell lines with a mutation in the transmembrane receptor NOTCH1 are responsive with high specificity. KRAS and, noteworthy, PIK3CA mutants remain resistant to these drugs.

The ability to uncover logic models around predefined operating points is an important requirement for practical application of such models. The implementation of additional constraints in the ILP formulation of LOBICO enabled us to systematically probe the ROC space and uncover logic models for different operating points of interest. To the best of our knowledge, LOBICO is the first method to provide this important capability.

## Discussion

Generally speaking, classification and regression approaches are primarily focused on prediction performance. Today’s state-of-the-art methods use complicated computational frameworks and data transformations to optimize how well the model fits the data. These models have little bearing on the mental model of the person using the approach. The interpretation of the model, if possible at all, is typically limited to a ranked list of important features.

Here, we have presented LOBICO, which, although also optimizing data fit, was developed specifically to produce models that are intuitively understandable. LOBICO generates small and robust logic models of binary input features that explain a continuous output. These models, which fit with standard formal reasoning, allow a researcher to easily assimilate the model with his or her domain. We maintain that the use of interpretable models is crucial in any scientific discipline, where researchers, not machines, generate hypotheses, gain novel insights and decide on further experimentation.

We have demonstrated LOBICO on a large cancer cell line panel by linking logic combinations of gene mutations to drug response. These logic models are easily integrated into current thinking based around stratifying patient populations using individual and combinations of gene mutations. LOBICO is, however, a general framework that can be applied to all research questions that can be described as a mapping from binary input features to a continuous output variable. For example, we successfully applied LOBICO to a cross of two natural yeast strains, the offspring of which were phenotyped for sporulation efficiency (**Supplementary Note 2** and Supplementary Figure 4). Importantly, although most (biological) measurements are not binary, they are often amenable to binarization or, at least, a binary interpretation. The use of binary variables allows for standard formal reasoning. That is, if the relation between variables is described using logic, it can be easily understood and reasoned with. We envision that LOBICO has many applications, both within biology as well as in other domains.

There are certain considerations about the applicability of LOBICO. First, LOBICO solves an NP-hard problem (**Supplementary Note 3**). We observed that the ILP solver can quickly traverse the search space of logic combinations when searching for small logic models. When looking for large models, on the other hand, the search space quickly explodes and it becomes prohibitive to find an optimal solution. However, we generally restrict LOBICO from inferring large models as they tend to overfit and are less interpretable.

We note that logic regression (LR) implements another search strategy based on simulated annealing, which also achieved a good performance on our dataset (**Supplementary Note 4**). In comparison, LR cannot guarantee that the obtained solution is optimal nor can it incorporate statistical performance constraints, which is the most relevant scenario for practical applications.

Second, the number of features and correlation structure within the feature space have a large impact on the feasibility of finding an optimal solution within a reasonable time. Large numbers of features drastically increase the search space. Hence, feature selection may be a prudent preprocessing step. When preselecting features, it would no longer be guaranteed that the optimal logic model is found. However, experiments on the cancer cell line panel show that features employed by the optimal logic model have high importance scores both in linear and non-linear regression models (**Supplementary Note 5** and Supplementary Figure 5). Thus, these methods might be useful for feature selection when LOBICO is to be applied to datasets with a prohibitively large number of features. Alternatively, expert knowledge can be used to preselect a smaller set of features. An advantage of the latter approach is that these features make sense to the domain experts and will lead to more easily interpretable models.

Correlated features are problematic, not only for LOBICO, but for most (sparse) linear or non-linear classification and regression models. Highly correlated features are effectively interchangeable. In practice, this means that only one feature is selected or that the importance in the model is spread across these features. An additional disadvantage for LOBICO is that the ILP solver will spend a long time deciding which of the correlated features should be part of the optimal solution. Also, from the perspective of interpretation, correlated features can lead to ambiguity and should preferably be dealt with prior to performing a LOBICO analysis.

The implementation of LOBICO as an ILP framework offers substantial advantages and possibilities. Using sophisticated ILP solvers, LOBICO traverses the huge search space of logic combinations fast enough for application to large datasets involving multiple model complexities and cross-validation. An argument against logical analysis is the loss of information when binarizing continuous data. The ILP formulation allows LOBICO to retain the continuous information in the form of sample-specific weights, which, as an added benefit, leads to robust models. Further, constraints guaranteeing predefined statistical performance criteria can be easily incorporated into the ILP framework, giving the logic models direct practical applicability.

A straightforward extension includes adjusting the ILP formulation to prioritize or even require certain features to be part of the inferred logic model. Also, certain combinations of features can be allowed or disallowed in the model. In this way, biological networks can be captured by ILP constraints, possibly reducing the search space and leading to logic models that reflect network biology. These ideas will be explored in future research.

## Methods and materials

### Cancer cell line dataset

We used a slightly expanded version of a previously published cancer cell line dataset [10]. Specifically, our dataset contained 714 cell lines and 142 drugs instead of 639 and 130, respectively, in the original publication. Although capillary sequencing was performed for 64 genes, our dataset only included the 54 genes for which at least one mutation or copy number aberration was found. Additionally, we included 6 known oncogenic gene fusions, resulting in 60 mutation features. The drug screening dataset is incomplete, i.e. not all 142 drugs have been screened across all 714 cell lines. In total 81,700 IC50s were measured and 19,688 (19%) were missing values. For the large majority of drugs, IC50s were obtained for between 600 to 700 cell lines. Since we applied LOBICO to each drug separately, cell lines that lack an IC50 were not used. We did not impute missing IC50s. In this work, IC50s were recorded as the natural logarithm of the half-maximal inhibitory μM concentration. Complete data and information on cell lines, the binary mutation matrix, drugs and IC50s are found in **Supplementary Data 2** and **4**. Additional information on these data is found in [10, 17] and on the Genomics of Drugs Sensitivity in Cancer webpages (http://www.cancerrxgene.org/).

### Binarization thresholds of the IC50s

The binarization threshold for each of the drugs was automatically determined using a heuristic outlier procedure, which consists of four steps:

1. *Upsampling*. For a drug, we gathered the IC50s and their confidence intervals of all ( *n*) cell lines that were screened for that drug. Then, for each cell line, we took the IC50 and the confidence interval and used this to define a normal distribution. Specifically, the mean of this normal distribution was the IC50 and the standard deviation was the average difference between the IC50 and the lower and upper bound of the confidence interval. We sampled 1,000 data points from this normal distribution and repeated that for all cell lines, leading to 1,000·*n* data points in total.
2. *Density estimation*. We performed kernel density estimation on the 1,000· *n* data points using as kernel a normal distribution with bandwidth (standard deviation) 0. 5. This kernel density estimate, *f*, is defined on the interval from the minimum value of the 1,000·*n* data points, *f*^min^, to the maximum value, *f*^max^, and was normalized such that 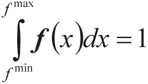.
3. *Modeling the population of resistant cell lines*. The population of resistant cell lines was modeled as a normal distribution. We used the mode (highest point) in *f* as the mean, *μ*, of this distribution. This choice was based on the expectation and previous observation that the large majority of cell lines is resistant to a drug [10]. To compute the standard deviation of this distribution, *σ*, we first computed the parameter *θ*, which marks the divide between sensitive and resistant cell lines. *Θ* was computed using the following rules. i. *Θ* is the maximum value, where the derivative *f*′ is zero, i.e. *f*′(*θ*) = 0, under the constraints that *θ* < *μ*, *f* (*θ*)< 0.8 · *f* (*μ*) and 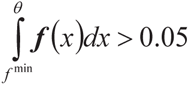.
ii. If no such value exists, *Θ* is the maximum value, where the second derivative *f*″ is zero, i.e. *f*″ (*Θ*) = 0, as well as *f*′″ (*θ*)> 0, again under the constraints that *θ* < *μ*, *f* (*θ*)< 0.8 *· f* (*μ*) and 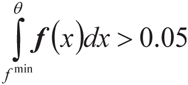.
iii. If no such value exists, *θ* was set to *f*^min^. If *θ* is found under rule i., this indicates bimodality in *f*, i.e. there is another peak in the distribution to the left of *μ*, which represents the distribution of sensitive cell lines. Similarly, if *θ* is found under rule ii., there is a marked change in the slope of *f*, which points to the distribution of sensitive cell lines. The standard deviation, *σ*, is computed as the median distance of all data points within the interval [*θ μ*]from *μ*. Finally, the population of resistant cell lines is represented by the Normal distribution *N*(*μ,σ*^2^),which we denote by *g*.
4. *Evaluating the cumulative Normal distribution to find the binarization threshold*. The binarization threshold, *b*, is controlled by parameter *t*. Specifically, *b* is chosen such that the cumulative Normal distribution function at *b* equals *t*, i.e. 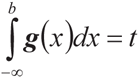. The default setting for *t* is 0.05. Cell lines with IC50s smaller than *b* are called sensitive, whereas cell lines with IC50s larger than or equal to *b* are called resistant.

Supplementary Figures 6, 7 and 8 provide a visual description of the four-step-procedure to binarize IC50s for three different drugs, where rule i., ii., and iii. were used to find *Θ*, respectively. Binarization thresholds (for *t* = 0.05) are found in **Supplementary Data 2**.

### LOBICO

The goal of LOBICO (Logic Optimization for Binary Input to Continuous Output) is to construct a Boolean logic function that optimally maps the binary variables of an input dataset to a continuous output variable. LOBICO is an extension to the work of Kamath *et al*.[18], which describes a way to solve the Boolean Function Synthesis Problem (BFSP) using integer linear programming (ILP). See **Supplementary Note 3** for a description of the BFSP. Similarly to the BFSP, LOBICO is an NP-hard problem (also in **Supplementary Note 3**).

LOBICO has three main inputs:

1. X, a *N* × *P* binary matrix with *N* samples characterized by *P* binary features *x*_1_,*x*_2_, …,*x_P_*. The *P* columns of X are denoted as **x**_1_,**x**_2_, …,**x**_*P*_.
2. y, a *N* × 1 binary vector, which is the binarized version of the continuous output variable.
3. w, a *N* × 1 continuous vector with weights for each of the *N* samples.

LOBICO finds an optimal logic function 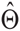 that minimizes the weighted sum of incorrectly inferred samples:

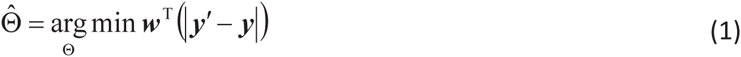

where, y′ = ∀(X) is a binary vector with the inferred binary labels. The logic functions inferred by LOBICO are in disjunctive normal form (DNF), a generalized logic notation also known as the sum-of-products expression. The complexity of a DNF is determined by two parameters: *K*, the number of disjunctive terms and *M*, the maximum number of selected features per disjunctive term.

#### IP formulation of LOBICO

The ILP formulation to find an optimal 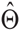 given X, y, w, *K* and *M* employs three variables: First, selection variables are introduced to determine which variable is part of a disjunctive term.

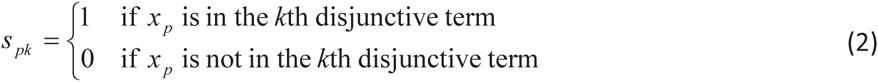

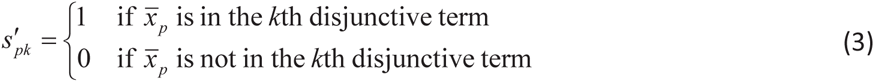

Second, auxiliary variables **t**_1_, …,**t**_*K*_, which are vectors of length *N*, are used to represent the disjunctive terms. Third, the disjunctive terms are combined in the final disjunction resulting in the inferred binary output variable y′. Figure 5 presents a graphical overview of the variables used in the ILP formulation.

**Figure 5.**
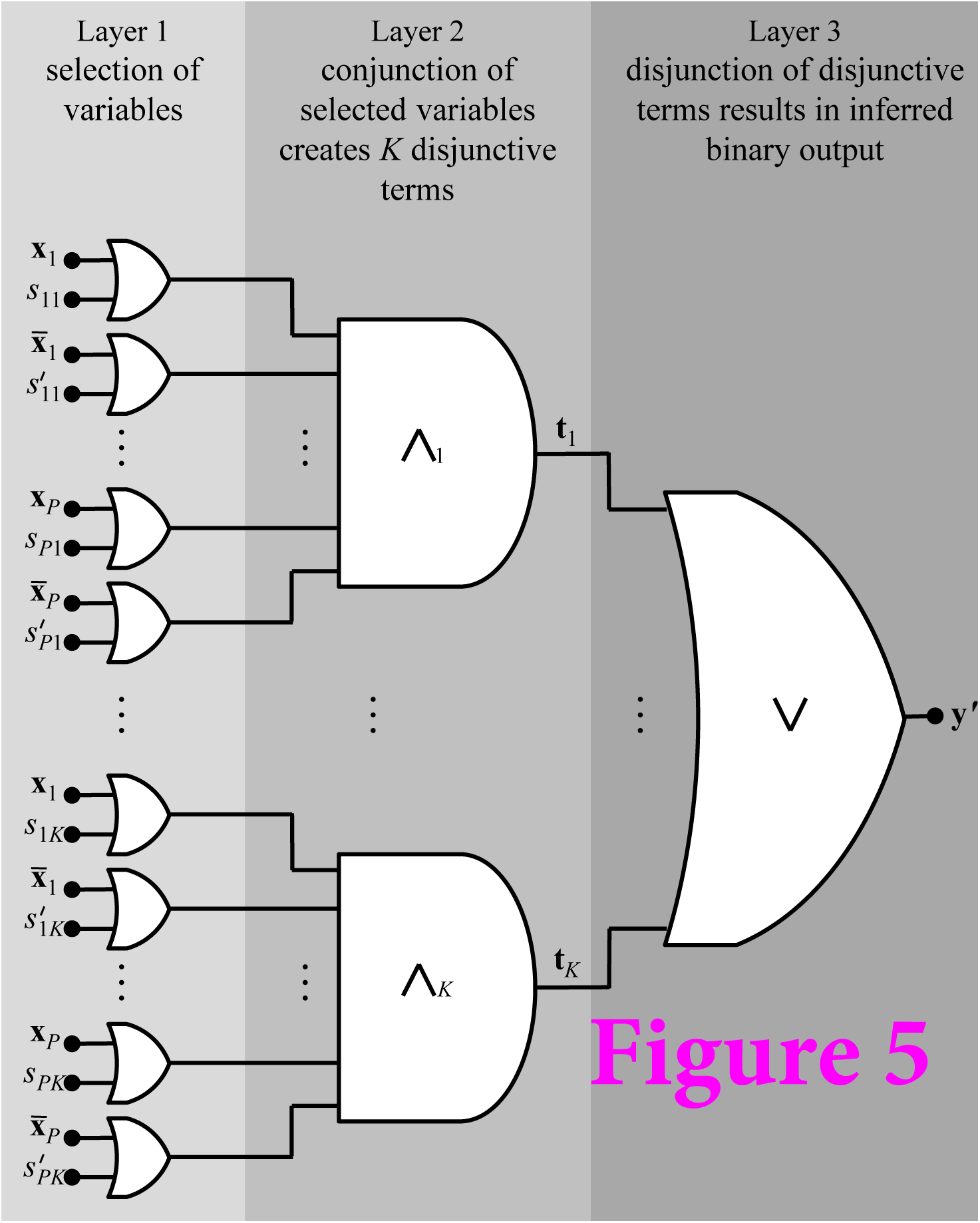
| 3-layer Boolean circuit representing the structure of the LOBICO ILP formulation. In Layer 1 variables s_11_..,*s_PK_* are used to select the inputs ( *x_1_,x_2_, …,x_P_*) that are combined using a conjunction (AND gate) to create the *K* disjunctive terms in Layer 2. These disjunctive terms (the outputs of the AND gates) are represented by variablest **t**_1_..,**t***_K_*. In Layer 3 the disjunctive terms are combined using a disjunction (OR gate) resulting in the inferred binary output variable y’. This figure is adapted from Figure 2.1 in Kamath *et al*. [18].

The complete ILP formulation is given below.

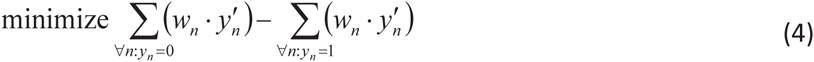

subject to

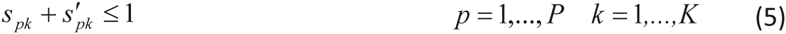

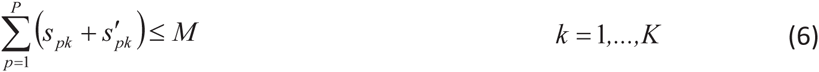

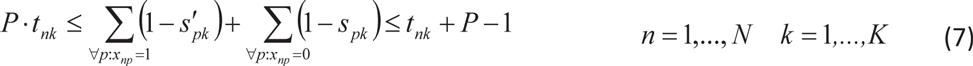

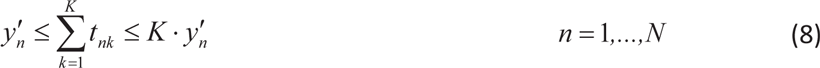

Following is a brief interpretation of the equations. The objective function in Equation 4 is the ILP formulation of Equation 1 and thus represents the weighted sum of incorrectly inferred samples, which should be minimized. The constraints in Equation 5 ensure that *x_p_* and its negation 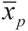 are not simultaneously part of the same disjunctive term. The constraints in Equation 6 ensure that the total number of selected features in a disjunctive term does not exceed *M*. Equation 7 encodes the AND-gates that define the *K* disjunctive terms. The output of the AND-gate *t_nk_* is only 1 for those samples, where the binary data in X agrees with all selected features, i.e. *x_np_* = 1 ∀*p*: *s_pk_* = 1, *x_np_* = 0 ∀*p*: *s*′*_pk_* = 1. In that case, the two summations in the middle part of the equation add up to *P*, thereby constraining *t_nk_* to 1. Equation 8 encodes the OR-gate, which combines the *K* disjunctive terms in the inferred binary output variable y′.

#### Feature importance scores

Feature importance (FI) scores are based on the activity measure of variables in Boolean networks [19, 20]. The importance score of feature *a* is defined as:

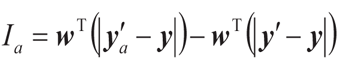

It represents the increase in the weighted sum of incorrectly inferred samples (Equation 1), henceforth called error, when comparing the optimal solution 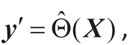, resulting in the error 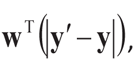,with 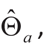, the optimal solution where feature *a* is left out of the model, resulting in the error 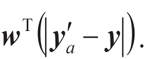.

In practice, leaving feature *a* out of the DNF is achieved by either setting all values of *a* to 1 in the case that feature *a* is part of a disjunctive term with at least one other feature, or setting all values of *a* to 0 in the case that feature *a* is the only feature in the disjunctive term. Features that are not part of the model (DNF) receive an importance score of 0.

See “Application to the cancer cell line panel” below, where we explain how we aggregated feature importance scores across suboptimal solutions, model complexities and CV folds to quantify the importance of gene mutations in explaining drug sensitivity.

#### Sensitivity and specificity constraints

The ILP formulation, which includes both the actual output (y) and inferred output (y’), enables the straightforward implementation of constraints, which guarantee that the ILP solution meets certain performance statistics (provided that a solution actually exists). Specifically, we implemented constraints on minimum sensitivity (also called true positive rate or recall), *TPR*^min^, and minimum specificity, (also called true negative rate or one minus the false positive rate), *TNR*^min^, as follows:

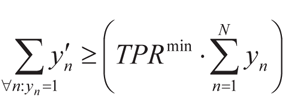

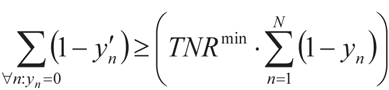

Note that in Equation 10 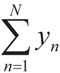 is the number of positives and 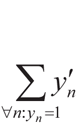 is the number of true positives. In Equation 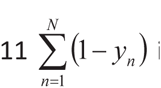 is the number of negatives and 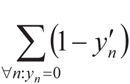 is the number of true negatives.

Using the notion of sample-specific weights (as represented by w), we defined ‘continuous’ versions of constraints on the sensitivity and specificity as follows:

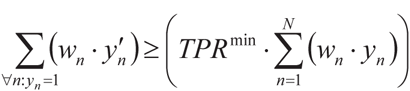

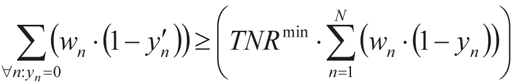

Visualizations of LOBICO solutions in the ROC space using the constraints on the ‘continuous’ sensitivity and specificity are found in **Supplementary Data 3**.

#### Application to the cancer cell line panel

***LOBICO inputs:*** LOBICO was applied to each drug in the cancer cell line panel separately. The three main inputs for each LOBICO analysis were:

1. X, the *N × P* binary mutation matrix with *N* cell lines and *P* = 60 binary gene mutations. *N* differs per drug as not all drugs have been screened across all cell lines. For the large majority of drugs, *N* is between 600 and 700.
2. y, the *N* x1 binary vector indicating whether a cell line is sensitive (1) or resistant (0) to the drug. This vector is obtained by binarizing the continuous IC50s using the binarization threshold, which is determined as explained above.
3. **w**, a *N*×1 continuous vector with weights for each of the *N* cell lines. w is simply the absolute difference between the IC50s and the binarization threshold, normalized per class:

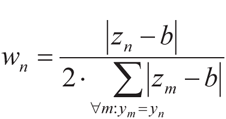

Here, *z_n_* is the continuous IC50 of cell line *n* and *b* the binarization threshold for the drug. The normalization ensures that both classes (sensitive and resistant) have the same total weight, i.e. 0.5. Note that 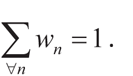.In “Use of continuous output yields more robust and accurate models”, where we compared to the setup in which we did not use the sample-specific weights, 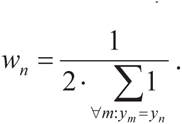.

***Statistical performance constraints:*** For the analyses described in “LOBICO finds logic models around a user-defined operating point”, we systematically varied *TPR*^mm^ (Equation 10) and *TNR*^min^ (Equation 11). For all other analyses in the paper, we did not employ the statistical performance constraints.

***Model complexities:*** We applied LOBICO with eight different DNF complexities, i.e. we used all combinations of natural numbers *K* and *M* provided that *K* ·*M*≤4 (see **Table 1**).

***Cross-validation:*** For each LOBICO analysis in the paper, a stratified 10-fold crossvalidation (CV) strategy was employed.

***Feature importance scores:*** We computed two types of FI scores for the analysis of the drug response models:

1. *Model-complexity-specific FI scores*. The model-complexity-specific (MCS) importance of feature *f* is the non-zero average of the FI scores of feature *f*, *I_f_* (Equation 9), obtained from the optimal solution and suboptimal logic models (if any) for particular model complexity (*K* and *M*). Often the suboptimal solutions differ only slightly in error with respect to the optimal model, but contain different features. (See “Solving the ILP using CPLEX” later in this section that explains how suboptimal solutions are found.) This means that the MCS FI score of feature *f* is the mean FI of *f* across all “good” logic models that contain feature *f*. Features that are neither part of the optimal logic model nor part of any suboptimal logic model get a score of zero. In this work, the sample-specific weights were normalized such that errors are between 0 and 1, with random prediction resulting in an expected error of 0.5 (Equation 14). Hence, FI scores are between 0 and 0.5, although the large majority (about 90%) of the non-zero FI scores are smaller than 0.05, and only a small portion (about 5%) are larger than 0.1. The MCS FI scores were used in the robustness analysis (Figure 3b and Supplementary Figure 2) and the ROC analysis (Figure 4). They are displayed in **Supplementary Data 1** and the upper part of Figure 3a.
2. *Aggregated FI scores*. The aggregated importance of feature *f* is the non-zero average of the MCS FI scores of all model complexities that have a CV error equal or smaller than the CV error for the single-predictor model ( *K = 1* and *M = 1*). Calculating the aggregated FI score involves averaging across the logic models based on all samples as well as the logic models based on the CV training folds. Hence, the aggregated FI scores provide a more comprehensive view of the importance of features in explaining the variation in the output variable. The aggregated FI scores were used in the ground truth analysis (Figure 3c). They are displayed in **Supplementary Data 2** and the lower part of Figure 3.

#### Solving the ILP using CPLEX

The ILP problems were solved using IBM ILOG CPLEX Optimization Studio V12.4, which is freely available for academic use. Importantly, ILP solvers guarantee optimal solutions (within a numerically small tolerance). Besides the optimal solution, we collected suboptimal solutions using CPLEX’s solution pool. Specifically, up to 25 solutions with a relative gap smaller than 0.1 were gathered using the default pool replacement strategy of replacing the least diverse solutions. We employed a time limit of 1 hour (3600 seconds) per ILP. Less than 1% of the ILP runs did not find the guaranteed optimal solution within this time limit (See Supplementary Figure 9). These solutions were all for 2x2 models. In these cases we used the best solutions found thus far. All other parameters were set to their default values.

#### LOBICO code availability

LOBICO is implemented in MATLAB and Python and is available through https://github.com/tkniinen/LOBICO.

## Supplementary Information

### Supplementary Note 1- Robustness across CVfolds

We investigated the robustness of the logic models across the ten CV training folds for each of the 142 drugs. The logic models for a drug were inferred using the model complexity (defined by *K* and *M*) selected by CV for that drug in the standard setting, i.e. with the sample-specific weights and t=0.05. The use of the continuous output resulted in a smaller variation in the FI scores across the CV folds (Supplementary Figure 2a). Particularly, the FI scores for the logic models across the CV folds had an average Pearson correlation coefficient larger than 0.75 for 113 drugs (80%), and 40 drugs (28%) had a correlation larger than 0.95. In contrast, for the logic models based on binarized data, there where 89 drugs (63%) had a correlation larger than 0.75 and 35 (25%) had a correlation larger than 0.95.

In comparison to changing the binarization threshold (Figure 3a), we observed that logic models inferred from the randomly sampled subsets, i.e. the CV folds, showed more variability in the FI scores, and thus a smaller correlation amongst the CV folds. We hypothesized that the inclusion or exclusion of samples, especially those far away from the binarization threshold, can have a large effect on the optimization function (Equation 1), and therefore a large effect on the inferred optimal logic model and the resulting FI scores. To test this hypothesis, we compared the similarity between CV folds with the similarity of the FI scores derived from the logic models trained on these CV folds. Specifically, for each of the 142 drugs separately, we computed:

1. for each pair of the CV training folds, say *a* and *b*, the similarity between the CV folds *a* and *b* in the following manner:

a. We took **w**, the *N*× 1 continuous vector with weights for each of the *N* samples. **w** is the absolute difference between the IC50s and the binarization threshold, normalized per class (Equation 14).
b. We created **w***^a^* and **w**^b^, where **w***^a^* is identical to **w**, except that all samples that are not part of the training set of *a* are replaced by 0, and similarly for **w**^b^.
c. As a metric of the similarility between CV folds *a* and *b*, we computed the Pearson correlation coefficient between vectors **w***^a^* and w^b^.
2. for each pair of the CV training folds, as a metric of the similarility of the FI scores between the two members *a* and *b*, the Pearson correlation coefficient between the FI score vectors derived from the logic models trained on CV folds *a* and *b*.

With the 142 drugs and 10-fold CV strategy, this resulted in 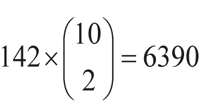 pairwise correlation scores for the similarity in the weight vectors and 6390 pairwise correlation scores for the similarity in the FI scores. We observed a clear relationship between these correlation scores (Supplementary Figure 2b). Particularly, pairs of CV folds with a small correlation between the weight vectors often had a small correlation between the FI scores. We observed that in about 5% of the cases the correlation between the weight vectors was quite low, i.e. the correlation coefficient was smaller than 0.7. These are cases, where the two CV folds include (and exclude) different samples with extreme IC50s, i.e. those far away from the binarization threshold. These ‘important’ samples have large weights in the weight vector, and when set to 0 in one of the folds, but not the other, lead to the low correlation scores between the folds. It is thus not surprising that the logic models inferred on these distinct CV folds lead to different FI scores.

This analysis confirmed our hypothesis that the larger variation in FI scores observed across CV folds is due to the inclusion or exclusion of samples with large weights, i.e. those far away from the binarization threshold.

### Supplementary Note 2- LOBICO on a yeast cross phenotyped for sporulation efficiency

We re-analyzed the genetic linking map of a cross of two natural yeast strains, a strain isolated from the bark of an oak tree that sporulates at 99% efficiency, and a strain originating from a wine barrel that sporulates at only 3.5% [21]. The genetic linkage map consists of 225 loci genotyped in 374 segregants. For each of the 374 recombinant offspring, the sporulation efficiency was measured as a percentage between 0 and 100. Gerke *et al*. [21] used composite interval mapping based on a stepwise regression model to find loci that significantly cosegregated with variation in sporulation efficiency, leading to 5 significant loci, L7-9, L10-14, L13-6, L7-17 and L11-2 (Table 1 in [21]). Next, a second stepwise regression was used to select significant predictors from the five loci and all 2 and 3-way interaction terms involving these five loci. The final model included three significant 2-way interaction effects and one 3-way interaction effect. All these interactions were comprised of combinations of the three most significant individual loci, i.e. L7-9, L10-14 and L13-6 (Table S2 in [21]).

We applied LOBICO to this dataset to evaluate which (logical) interaction effects would be uncovered. The genotype information was straightforwardly transformed into binary predictor variables: Alleles from the oak strain (wine strain) were set to 1 (0), resulting in a truth table with *n* = 225 loci and *p* = 374 segregants. The sporulation phenotype data was binarized by applying a threshold of 50%. Samples were weighted using the distance to this threshold. No specificity and sensitivity constraints were applied. We employed the eight model complexities also used for the cell line panel analysis, i.e. all combinations of *K* and *M* with *K.M* ≤ 4.

The largest single effect found in Gerke *et al*., loci L7-9, is also the best single predictor uncovered by LOBICO (Supplementary Figure 4). The two-input AND model found by LOBICO consisted of loci L7-9 and L10-14. This interaction, which is also one of the 2-way interaction effects found in Gerke *et al*. has a much higher specificity and precision than the single locus model, although a smaller recall. Many of the offspring with the highest sporulation efficiency have both the L7-9 and the L10-14 locus from the oak strain. The best model according to CV is a 2-by-2 model, which contains the same three loci as in the interaction effects found in Gerke *et al*., i.e. L7-9, L10-14 and L13-6. (Actually, the LOBICO 2- by-2 model contained L13-7 instead of L13-6; they are highly correlated. The fourth feature in the 2-by-2 is L7-11, which is highly correlated to L7-9.) Thus, LOBICO finds interactions between the same three loci as the regression model employed by Gerke *et al*..

It is important to point out that LOBICO uncovered these interactions using the complete dataset of 225 loci, and not by first filtering on individual features as was done in Gerke *et al*.. Surely, the (biological) interpretation of the logic model and the additive linear model is quite different. We would argue that the logic model is more intuitive and sensible than the linear model.

### Supplementary Note 3- Explanation of the Boolean Function Synthesis Problem and proof that LOBICO is NP-complete

The Boolean Function Synthesis Problem (BFSP) is a particular type of Boolean Satisfiability Problem, where the goal is to find an algebraic sum-of-products expression for an incompletely specified Boolean function Φ: {0,1}^n^ →; {0,l}. The sum-of-products expression is also called a disjunctive normal form (DNF), i.e. a disjunction of conjunctions. Each Boolean function can be expression in DNF. An element of the domain of Φ is called a minterm of Φ. The set of minterms for which Φ evaluates to 1 (resp. 0) is called the ON-set (resp.OFF-set). An incompletely specified Boolean function is one for which |ON-set| + |OFF-set| <2^n^. Supplementary Figure 10 displays an incompletely specified Boolean function with *n* = 10 input variables, *x*_1_,*x*_2_, …, *x*_10_ and an output variable *y*.

The number of rows in the Boolean truth table is given by *p* (*p* = |ON-set| + |OFF-set|), and is 40 in this case (40 << 2^10^). Note that in most biology applications, Φ is incompletely specified. The sought after algebraic expression is a Boolean DNF expression that evaluates to 1 for all minterms in the ON-set ( OFF-set) and to 0 for all minterms in the OFF-set. Formally, the problem is as follows: Given an ON-set and an OFF-set of minterms that characterize a Boolean function Φ, find a DNF of Φ with maximally *K* disjunctive terms having each maximally *M* variables. The corresponding decision problem is NP-complete [22].

The decision version of LOBICO is as follows: Given inputs X, y, w, *K*,*M* and a parameter *L*, does there exist a logic function 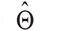 expressed in DNF(*K,M*), i.e. a DNF having at most *K* disjunctive terms and *M* literals, such that the weighted sum of incorrectly inferred samples as described in Equation 1 is less than or equal to *ε* ? Cleary, the problem is in NP. It is easily shown that this problem is also NP-complete by the following polynomial-time reduction from BFSP: Given an instance of BFSP we construct an instance for LOBICO by deriving **X** from the minterms, **y** from the ON-set and OFF-set and by setting **w** to **1**, i.e. *w_n_* = 1 *∀n*. Now, BFSP can be satisfied if and only if LOBICO has a solution with an error of *ε* = 0. Since the BFSP decision problem is NP-complete, the LOBICO decision problem is also NP-complete.

### Supplementary Note 4- Comparison with logic regression

Logic regression (LR) [3, 4] is a generalized regression methodology that can be applied to data with binary predictors, although continuous predictors are also allowed. The goal of LR is to find linearly weighted logic combinations of the original predictors that explain a continuous response variable or class label. We configured the implementation of LR, i.e. the R-package ‘logreg’, such that it infers logic models with a predefined model complexity. Specifically, logreg has a scoring function for classification using sample-specific weights, which we used to give it the same objective function as LOBICO (Equation 1). Also, logreg can be configured to output a single logic model (a tree) with predefined logical operators and size. (See below for experimental details.)

We ran LR for each of the 142 drugs in the cancer cell line panel using the model complexity (defined by *K* and *M*) selected by CV for the associated drug when using LOBICO. LR was run on the same computers (Intel(R) Xeon(R) CPU, E5645, 2.40GHz, 6 cores) as LOBICO and was given the same amount of CPU time (Supplementary Figure 11a). Then, we evaluated the logic formulas inferred by LR. Specifically, we looked at the Jaccard similarity of the selected predictors in the inferred LOBICO and LR models. (In computing the Jaccard similarity negated terms, e.g. -TP53, are treated as separate predictors from their positive equivalents.) For models with *K*=1 and/or *M*=1, a Jaccard similarity of 1 indicates that the exact same logic formula was found. For *K*=2 and **M**=2 (the 2×2 models) this is not necessarily the case, but we were not able to restrict logreg to output a DNF with *K*=2 and *M*=2 anyway. For example, for the drug ‘MG-132’ LOBICO inferred the 2x2 ‘(-MYC & RB1) | (-PIK3CA & -TP53)’, whereas LR inferred ‘(((-TP53) or (RB1 or NOTCH1)) and (-PIK3CA))’. The LR model is clearly not a DNF with *K*=2 and *M*=2.

Overall, LR found the same (optimal) logic formulas as LOBICO (Supplementary Figure 11b). The main exception is the 2×2 model (*K*=2, *M*=2), but this is because of the reason mentioned above. For the 4-input OR models (*K*=4, *M*=1) we observed four cases where the logic formulas differed between LOBICO and LR. Upon further inspection, we found that in these cases the formula inferred by LR had the same (optimal) error as the LOBICO solution. These LR solutions were present in LOBICO’s solution pool, i.e. they were part of the set of (sub-)optimal solutions output by LOBICO. (See experimental details below.)

In conclusion, when logreg parameters are properly set, LR can find the optimal solution when given the same amount of time that was necessary for LOBICO to find the optimal solution on the cancer cell line dataset. Potentially, LR finds this solution faster than LOBICO on this dataset. It is however important to point out that LR cannot guarantee that the obtained solution is optimal, and it is known that ILP solvers spend a long time proving that the found solution is indeed optimal. In future work, we will investigate the performance of LR and LOBICO on other (larger) datasets, and assess how the two methods can be used in parallel to find optimal solutions faster. For example, we will investigate whether LR can be used to identify initial starting models for LOBICO.

Importantly, LR cannot incorporate statistical performance constraints, such as sensitivity and specificity (Equations 10 - 13), which we assert is the preferred and, in practice, most relevant scenario for LOBICO inferences. Additionally, in contrast to LR, LOBICO can output the pool of (sub-)optimal solutions, which we used to measure feature importance.

Experimental details: LR was run using the R-packing logreg (version 1.5.8). We used simulated annealing as this search algorithm gave the best results. We followed the logreg’s documentation to set the upper and lower temperature of the annealing chain based on experiments with the cancer cell line panel. The number of iterations was set, such that the total CPU time spent on solving the problem was comparable to the CPU time that LOBICO needed to find the optimal solution (Supplementary Figure 11a). The simulated annealing parameters were set as follows (R-code):

> myanneal <- logreg.anneal.control(start = 1, end = -5, iter = T*200000, update = T*20000)

To infer a LR model with the same model complexity as LOBICO, we made sure that for 2-, 3- and 4-input AND models only AND operators were allowed. Similarly, for 2-, 3- and 4-input OR models only OR operators were allowed. For 2×2 models we allowed both AND and OR operators. The parameters of the logic ‘tree’ shape were set as follows (R-code):

> if (K>M) mytreecontrol <- logreg.tree.control(opers=3) else mytreecontrol <- logreg.tree.control(opers=2)
>
> if (K==2&M==2) mytreecontrol <- logreg.tree.control(opers=1)

LR was run to output one logic tree (ntrees=1), where the maximum number of leaves was set to *K*×*M* (nleaves=K*M). In the R-code below Y is y, X is **X** and W is **w** as used in the **Methods Section** and Equation 1. LR was run as follows (R-code):

> q<-logreg(resp=Y, bin=X, wgt=W, type=1, select=1, ntrees=1, nleaves=K*M,anneal.control = myanneal, tree.control = mytreecontrol)

### Supplementary Note 5- Comparison with sparse linear regression and Random Forests

We compared the LOBICO models obtained on the cancer cell line panel with Elastic Net [1], a sparse linear regression model, and with Random Forests regression [2], a nonlinear regression model. Specifically, for the 25 drugs with the lowest CV error in the original analysis, we compared the model-specific FI scores (of the model complexity selected by CV) with the regression weights inferred by Elastic Net (EN) and the importance scores inferred by Random Forests (RF).

We observed a large concordance between LOBICO’s FI scores and the EN regression weights (Supplementary Figure 5a). In the EN models, the large majority (63%) of all regression weights across the 60 features and 25 drugs were 0. Importantly, all of the important features according to LOBICO (FI>0.05) had a non-zero regression weight in EN. Moreover, the smallest EN regression weight for which the corresponding LOBICO FI was larger than 0.05, was 0.2752 (blue line in Supplementary Figure 5a), which was in the tail of the EN weights.

Similarly for RF, we observed a high degree of correlation between LOBICO’s FI scores and the RF importance scores (Supplementary Figure 5b). The important features according to LOBICO (FI>0.05) also had a high RF importance score. The smallest RF importance score for which the corresponding LOBICO FI was larger than 0.05, was 0.014, which marked the 82% percentile of the RF importance scores.

Experimental details: For EN, we used the MATLAB ‘lasso’ function with an alpha (mix between L2 and L1 penalty) of 0.5, 10-fold CV and sample weights w, the *N*xl continuous vector with weights for each of the *N* samples (Equation 14). For RF, we employed the Random Forests implementation for MATLAB v0.02 downloaded from http://code.google.com/p7randomforest-matlab/. The RF regression models were run with 1000 trees each and default settings for the other parameters were used. The reported importance scores represent the mean decrease in accuracy. To accommodate the different sample weights, we created (for each drug) a dataset of 10,000 samples, which were randomly drawn with replacement from the original dataset, where the probability of being drawn was proportional to the sample weights in **w**.

**Supplementary Figure 1.**
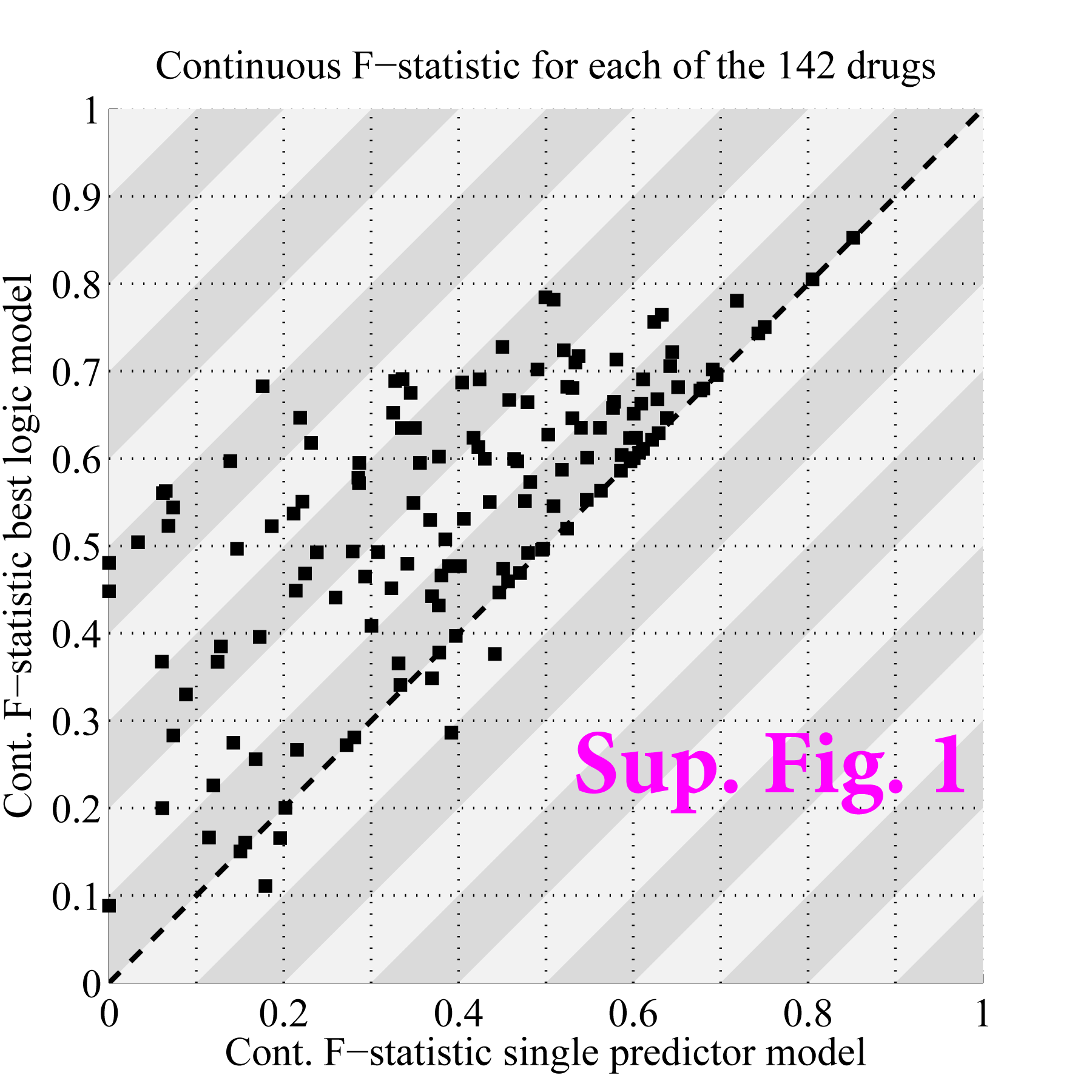
Multi-predictor models outperform single-predictor models. Scatter plot with the continuous F-statistic for single predictor models (x-axis) and the best (lowest CV error) multi-predictor model (y-axis). Each point represents one of the 142 drugs. The continuous F-statistic is the defined as the harmonic mean of the continuous recall and continuous precision. By analogy to Equations 12 and 13, the continuous recall and continuous precision are defined as 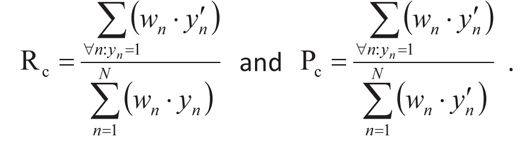. The continuous F-statistic uses the sample-specific weights that LOBICO uses in its optimization and is therefore a better performance measure than the standard F-statistic. Similarly, to the CV error depicted in Figure 2, the continuous F-statistic was computed on the inferred class labels of the samples in the test sets.

**Supplementary Figure 2.**
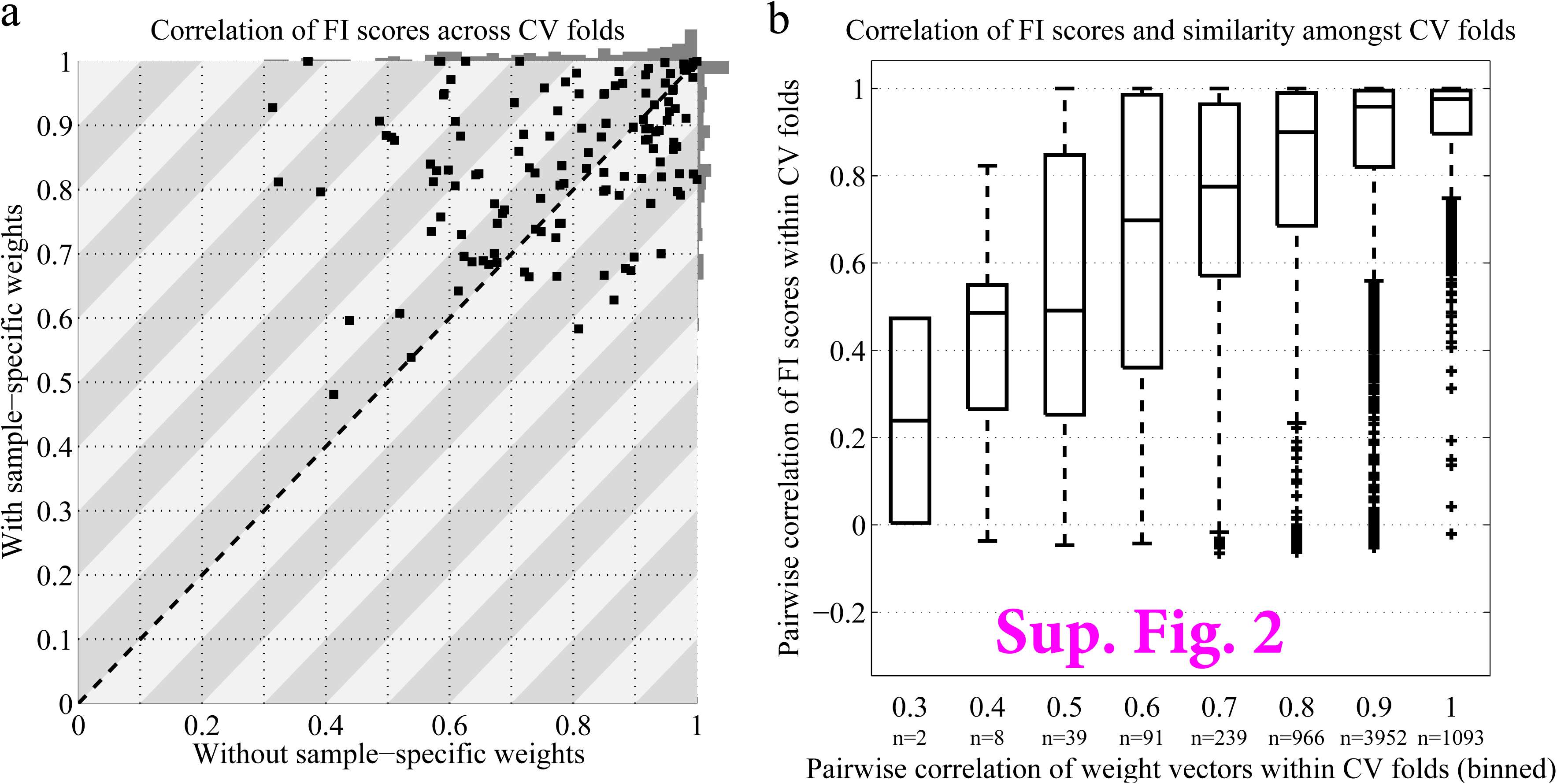
Robustness across CV folds. a) Scatter plot with the average Pearson correlation coefficients of the similarity of FI scores across the 10 CV folds for inferred logic models without (x-axis) and with (y-axis) the sample-specific weights. Each point represents one of the 142 drugs. The correlation scores are computed using the model-complexity-specific FI scores. The grey bars on top and to the right of the scatter plot represent histograms of these correlation scores for models without and with the sample-specific weights, respectively. **b)** Boxplot comparing the pairwise correlation of weight vectors between CV folds (x-axis) with the pairwise correlations of FI scores between the same CV folds (y-axis). The pairwise correlation of weight vectors were binned by rounding the correlation to the nearest decimal. The number of correlations per box is indicated below the box.

**Supplementary Figure 3.**
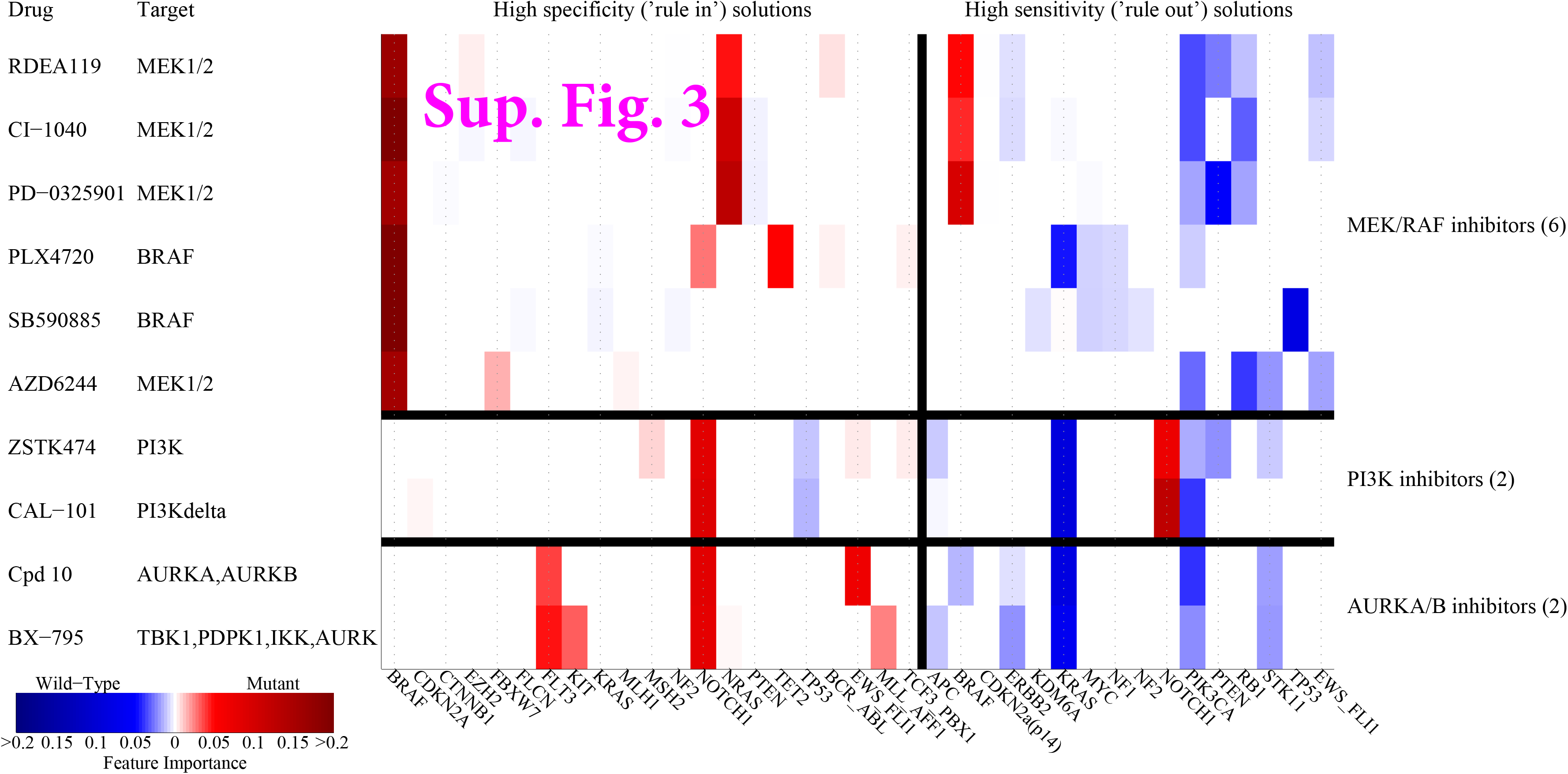
Feature importance scores for ‘rule in’ and ‘rule out’ solutions. FI scores of 6 MEK/RAF, 2 PI3K and 2 AURKA/B inhibitors (rows) for high specificity (‘rule in’) solutions (left) and high sensitivity (‘rule out’) solutions (right). High specificity solutions were defined as solutions with FPR<10%. Conversely, high sensitivity solutions were defined as solutions with TPR>90%. We distinguished between positive terms, indicating mutations (red) and negated terms, indicating wild-type (blue).

**Supplementary Figure 4.**
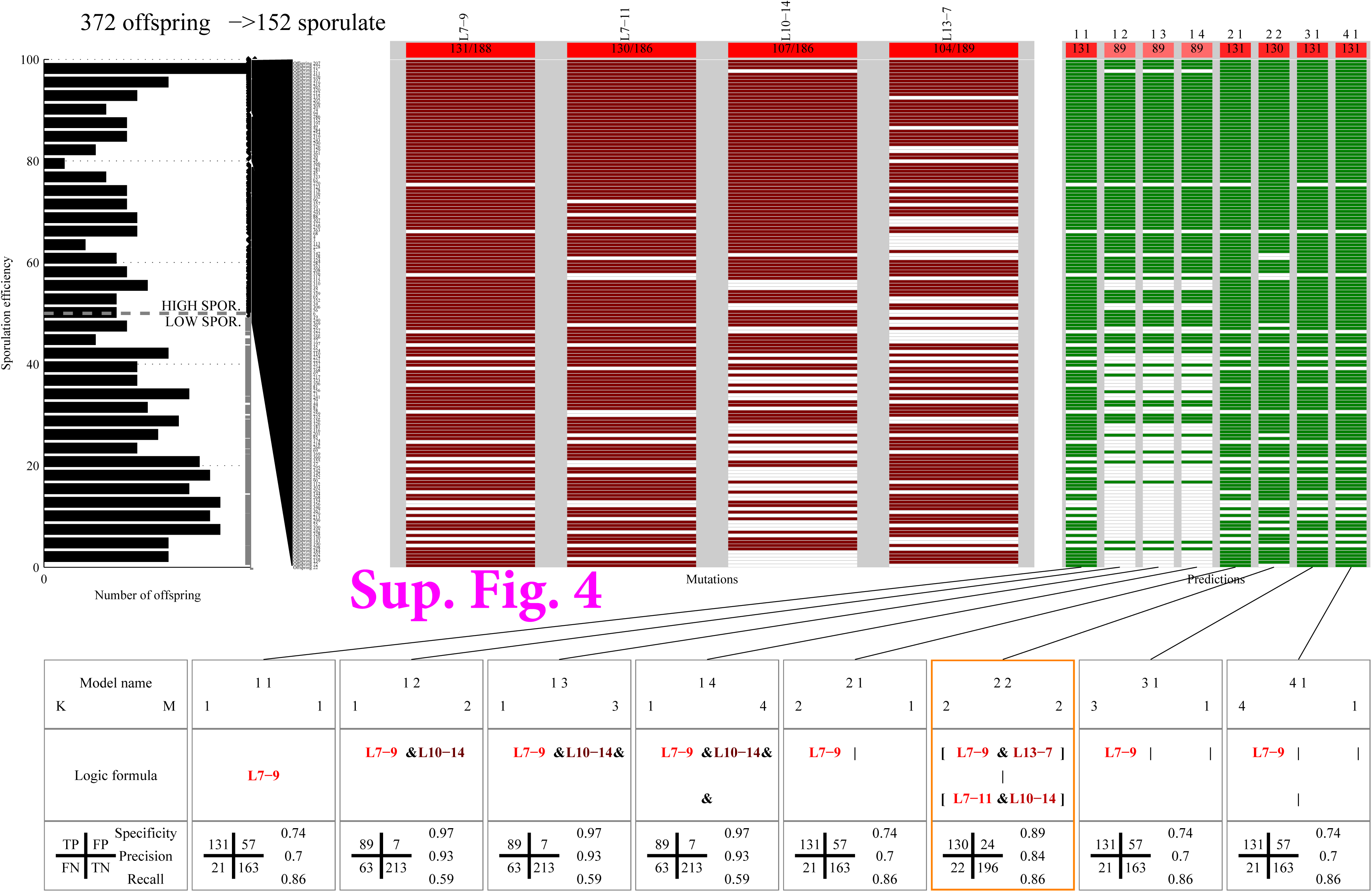
LOBICO results of the yeast cross phenotyped for sporulation efficiency. Standard LOBICO visualization of the uncovered logic models for the yeast cross dataset (Supplementary Note 2). See **Supplementary Data 1** for an explanation of the visualization.

**Supplementary Figure 5.**
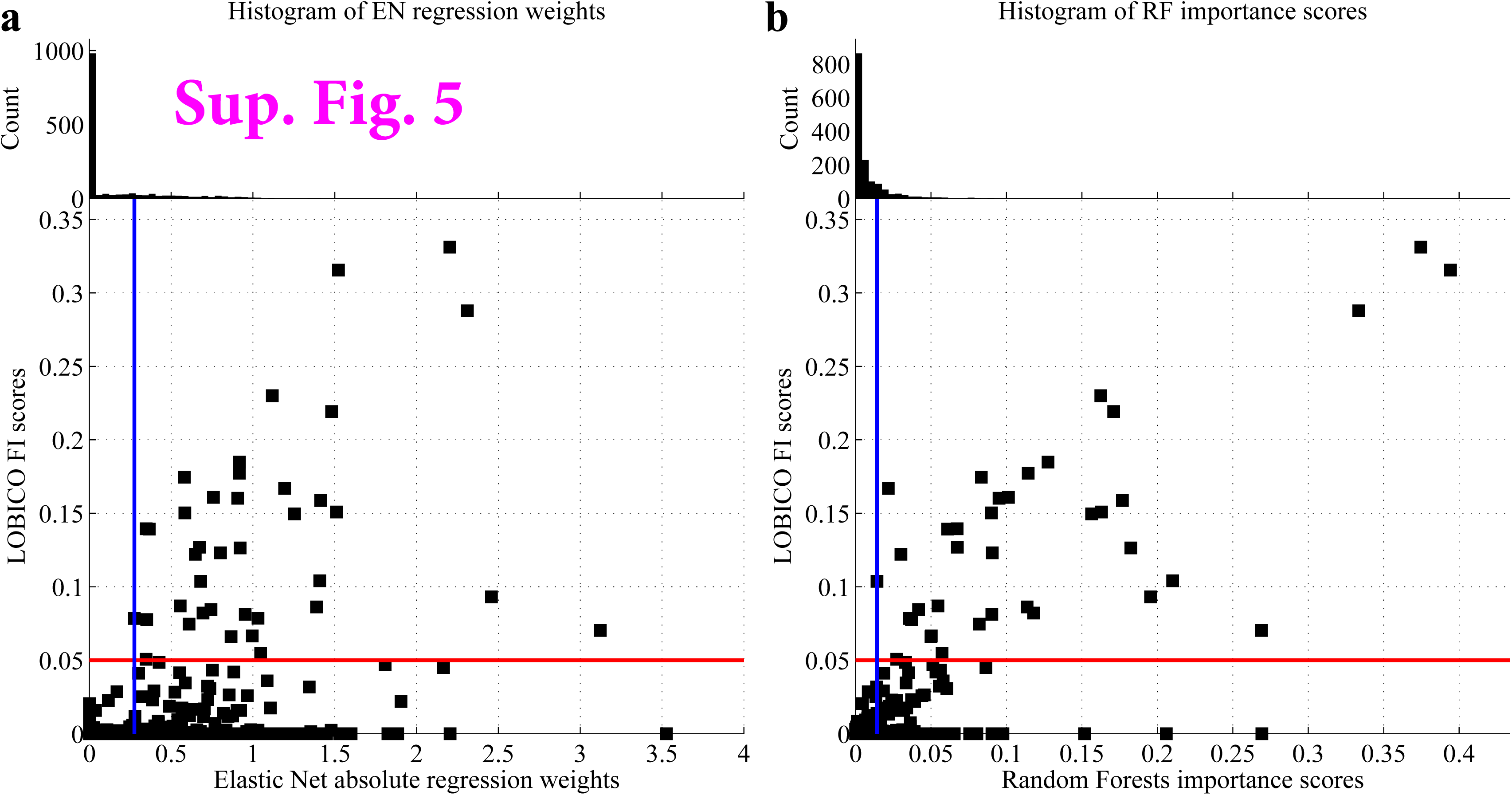
Comparison of feature importance scores between LOBICO, Elastic Net and Random Forests. a) Scatter plot comparing LOBICO’s FI scores with EN’s absolute regression weights. These scores and weights are derived from inferred models of the 25 drugs with the lowest CV error in the LOBICO analysis (Supplementary Note 5). The red line depicts the FI score of 0.05; features with a FI>0.05 are considered important predictors. The blue depicts the minimal EN regression weight for which the corresponding LOBICO FI was larger than 0.05. b) Similar to a), except LOBICO’s FI scores are compared to RF’s importance scores.

**Supplementary Figure 6.**
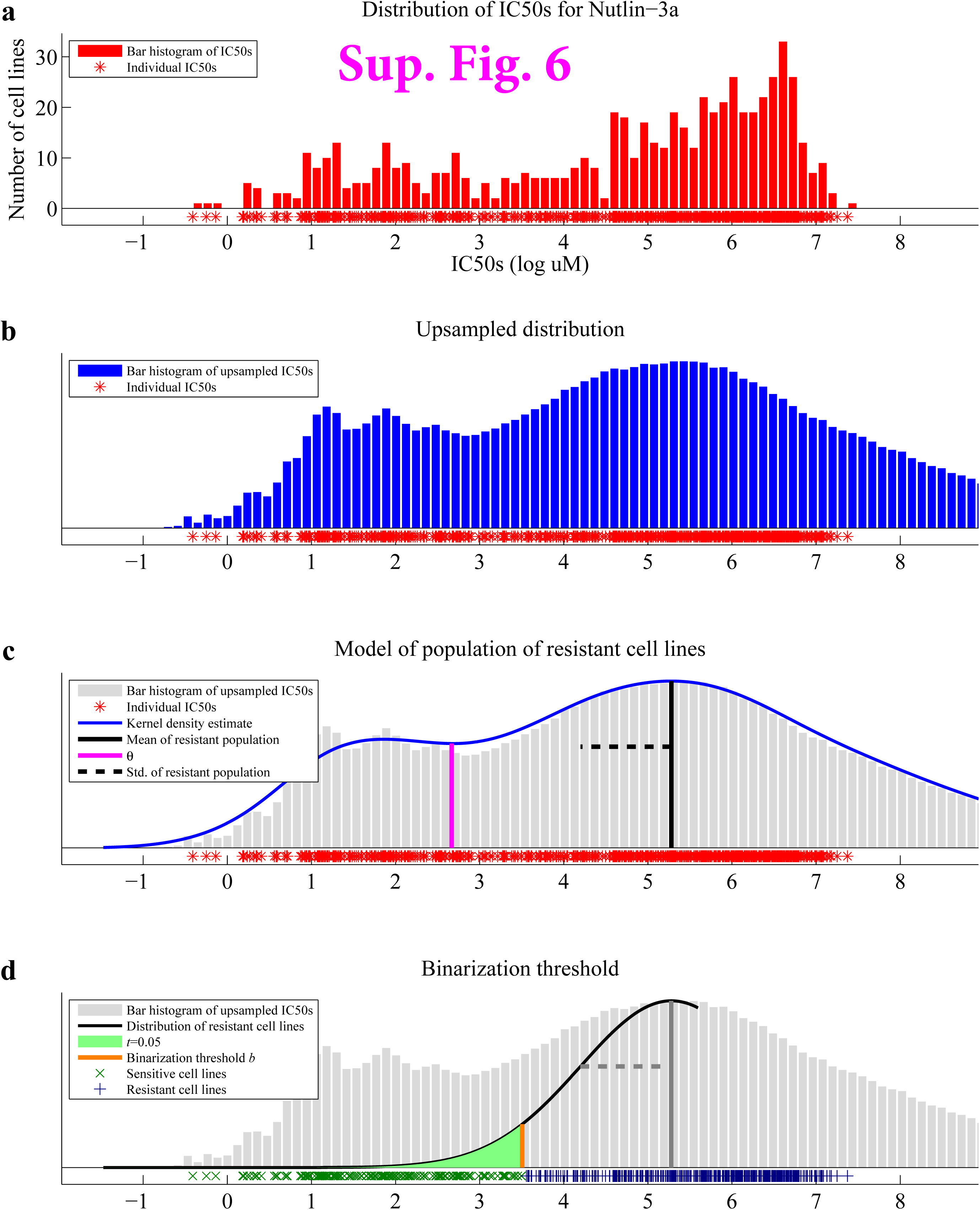
| Four-step-procedure to binarize IC50s for Nutlin-3a. **a)** Histogram plot for the distribution of IC50s for the drug Nutlin-3a. **b)** Histogram plot for the upsampled distribution **c)** Visualization of an empirical model (obtained through density estimation) of the upsampled IC50s (depicted in blue). *Θ* was computed using rule i. (See **Methods Section** for details.) **d)** Visualization of the model of resistant cell lines (depicted in black), from which the binarization threshold *b* (depicted in orange) is derived.

**Supplementary Figure 7.**
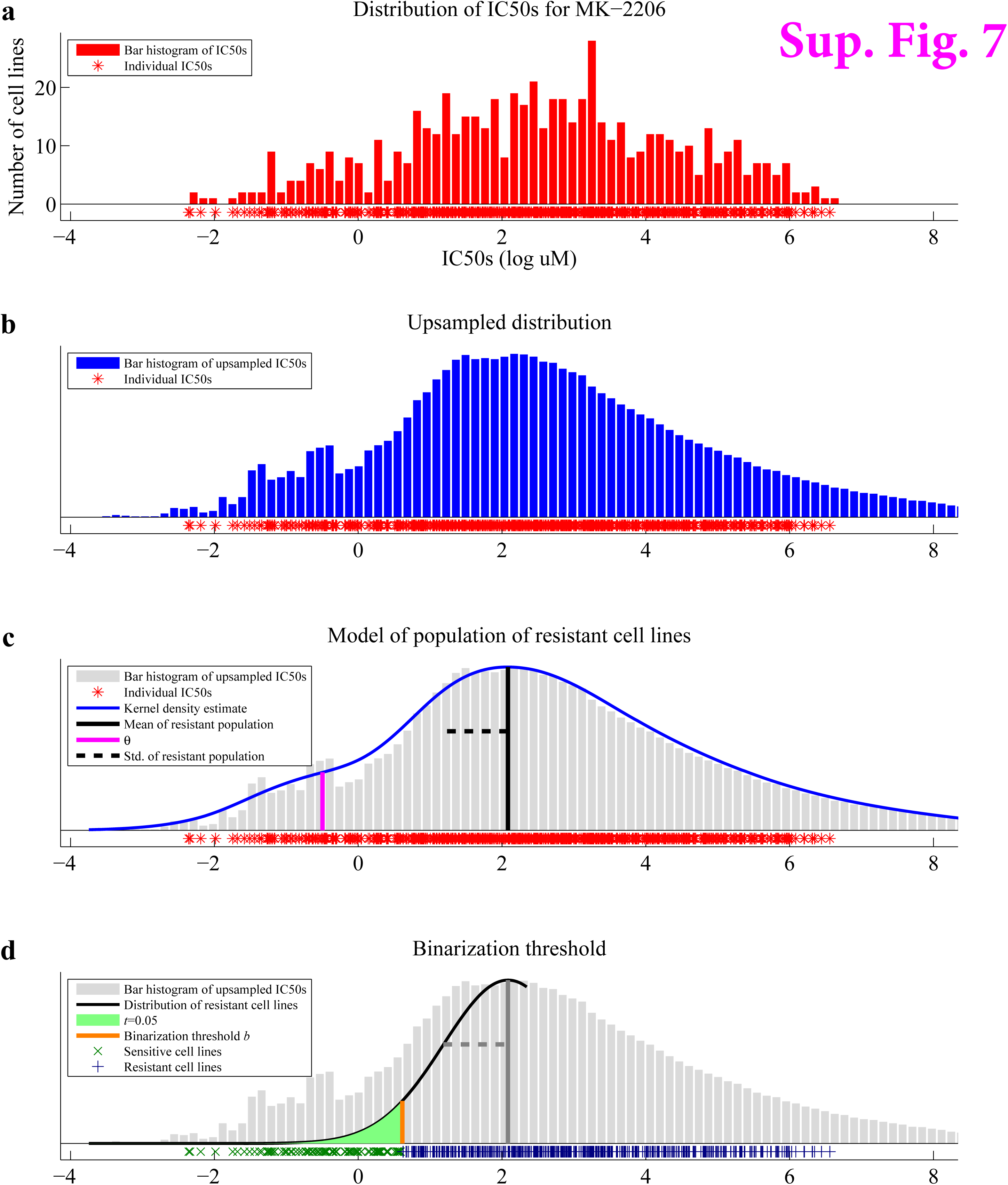
| Four-step-procedure to binarize IC50s for MK-2206. Similar to Supplementary Figure 6, except showing the procedure for drug MK-2206, and the use of rule ii to find *θ*.

**Supplementary Figure 8.**
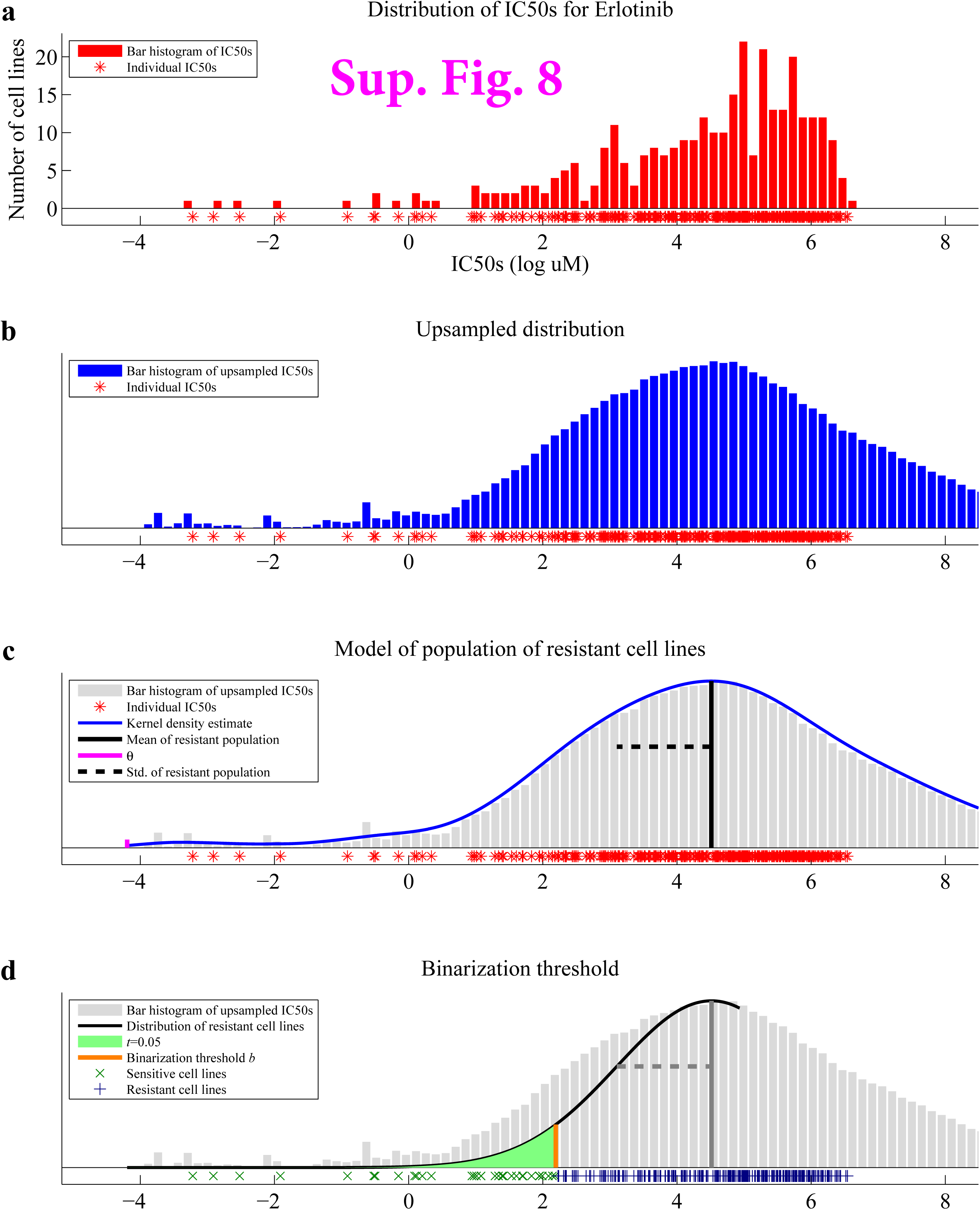
| Four-step-procedure to binarize IC50s for Erlotinib. Similar to Supplementary Figure 6, except showing the procedure for drug Erlotinib, and the use of rule iii to find *θ*.

**Supplementary Figure 9.**
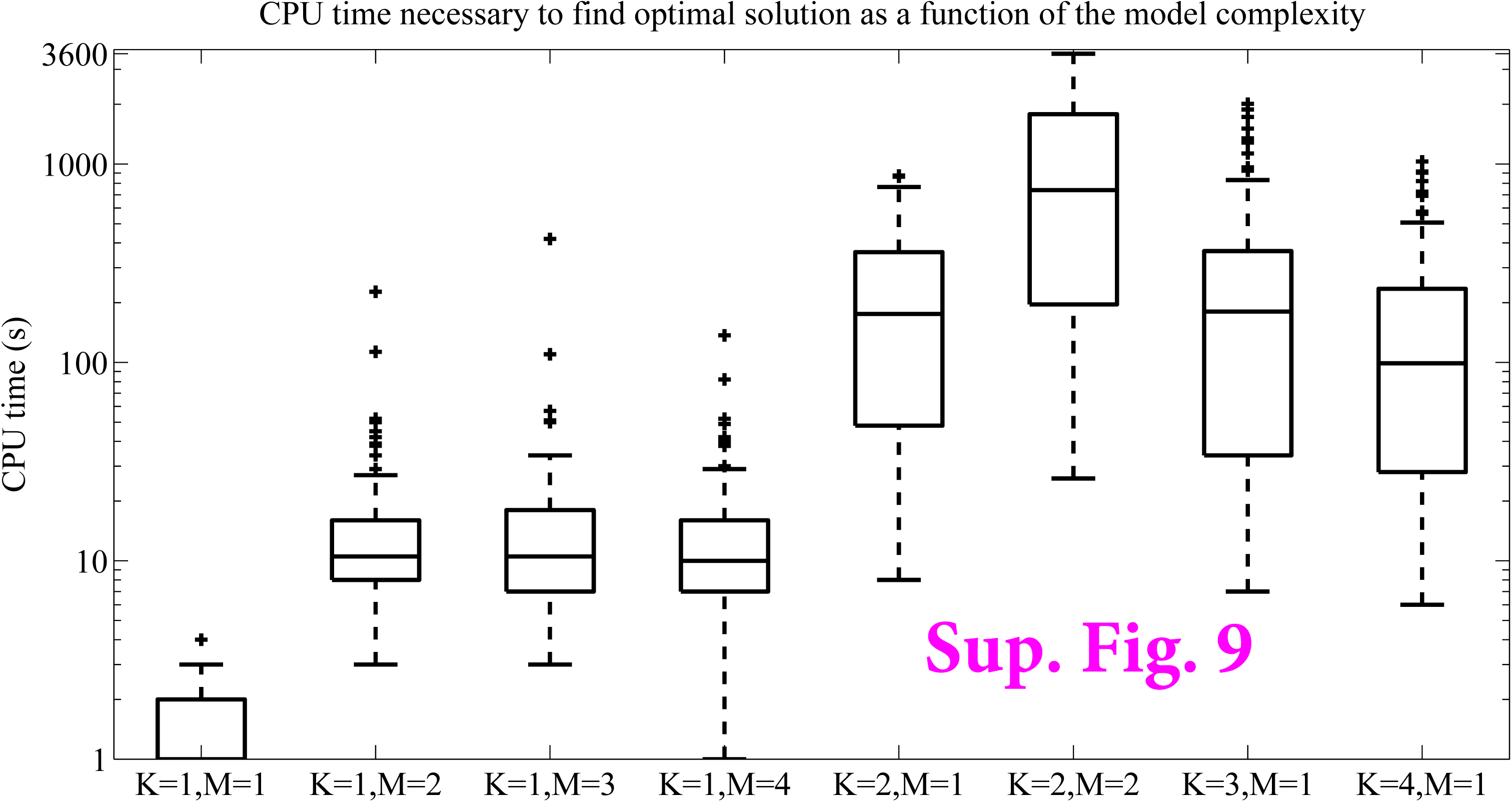
| Time needed to find optimal solution. Boxplot of CPU time (y-axis) necessary to find the optimal solution as a function of the model complexity (x-axis). Each box is comprised of 142 values, i.e. the time necessary for CPLEX to find the optimal solution with the indicated model complexity for each of the 142 drugs. These experiments were performed on a computer cluster, where each ILP was run on one node (Intel(R) Xeon(R) CPU, E5645, 2.40GHz, 6 cores) at a time.

**Supplementary Figure 10.**
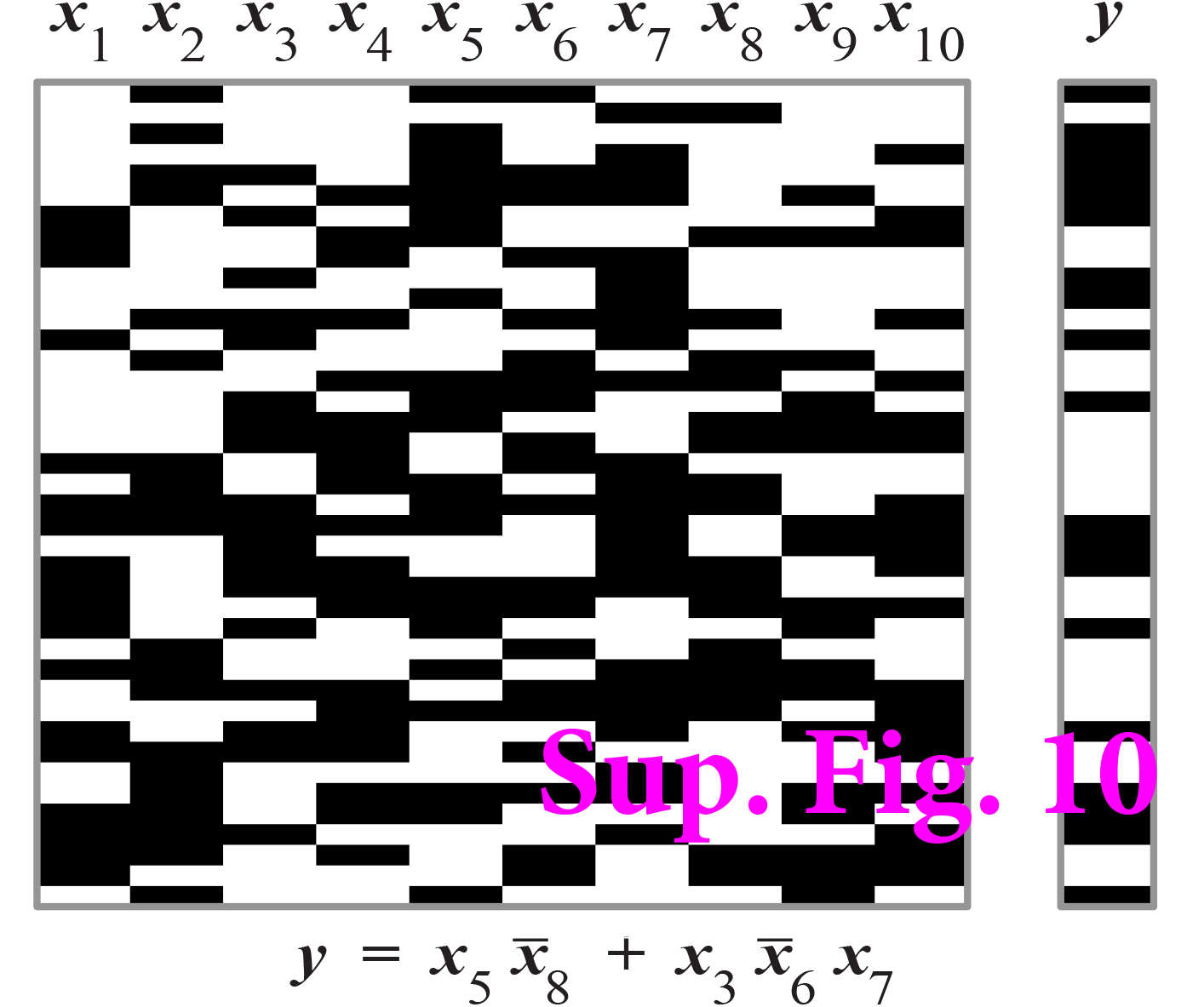
| Example Boolean truth table. Boolean truth table (black=1, white=0) with 10 input variables *x*_1_,*x*_2_, …, *x*_10_ and output variable *y*. The DNF expression with *K* = 2 and *M* = 3 for the truth table depicted is below the table.

**Supplementary Figure 11.**
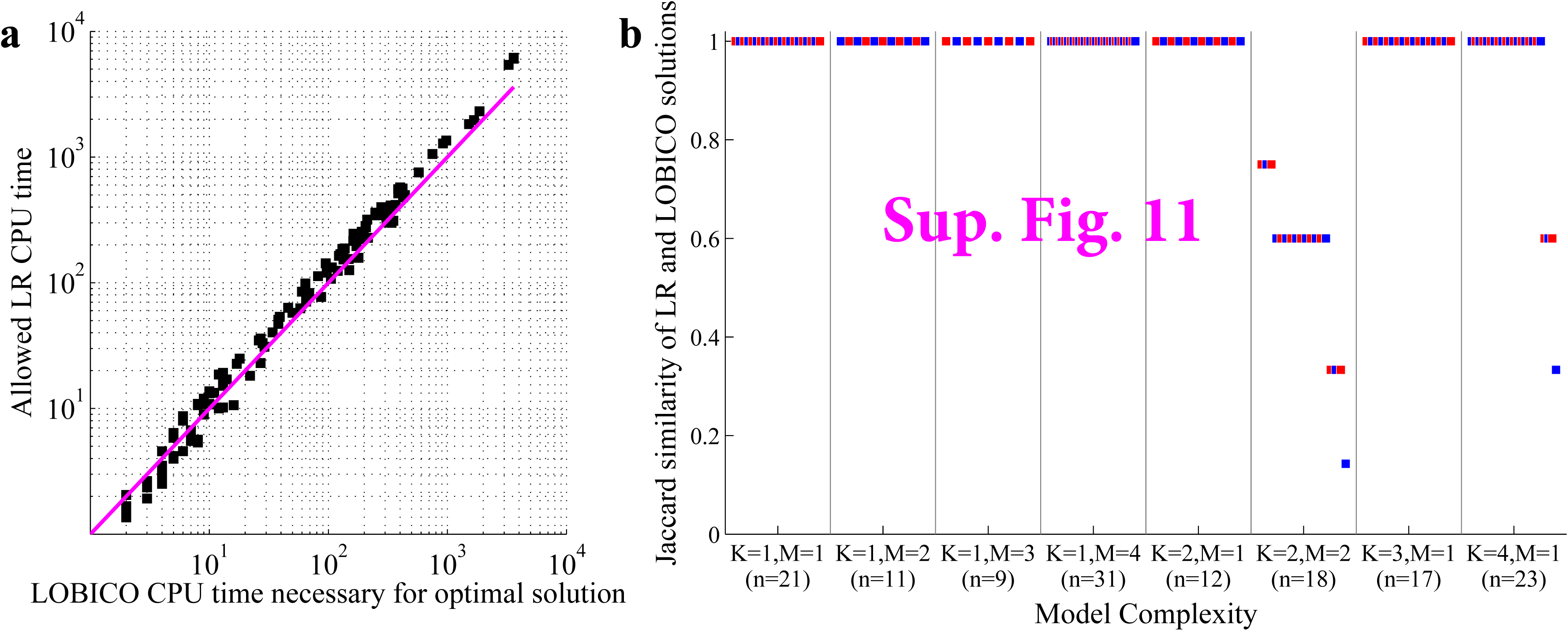
| Comparison of solution time and uncovered logic models between LOBICO and logic regression. **a)** Scatter plot comparing the CPU time needed for LOBICO to find the optimal solution (x-axis) and the CPU time given to LR to find the best solution (y-axis). Each of the 142 drugs is represented by a point. The magenta line is y=x. **b**) Plot of the Jaccard similarity between the LR and LOBICO solutions. Each of the 142 drugs is represented by a point. The points are alternately colored in blue and red for visibility. The drugs are grouped based on the model complexity (x-axis) for which the LOBICO and LR models were inferred.

## Supplementary Data Explanation

### Supplementary Data 1 | LOBICO visualization of the inferred logic models for all 142 drugs

PDF with a visualization of the LOBICO results for each drug (pages 8-149). The first 7 pages provide a visual explanation of the visualization.

### Supplementary Data 2 | Drug information

Tab separated Excel table containing information about the 142 drugs. Specifically, (from left to right in the table), information about the drugs (ID, name and target), binarization thresholds, ground truth mapping to the gene mutation features, model performance statistics of the inferred logic models, and aggregated feature importance scores for the gene mutation features in the inferred logic models.

### Supplementary Data 3 | 25 ROC models visual

PDF with visualizations of LOBICO solutions in the ROC space for each of 25 drugs with the lowest CV error in the original analysis. Blue crosses indicate the true positive rate (TPR) and false positive rate (FPR) at which the solution was found. The logic formula of the solutions is printed next to the blue crosses. The color of the genes in a formula indicate their FI. Colors range from black (moderately important) to bright red (highly important). For comparison, the best single-predictor solutions are visualized in green. The inlay depicts the histogram of IC50s of the drug together with the binarization threshold.

The PDF consists of 50 pages; each of the 25 drugs is represented by two visualizations: one using the standard (binary) definitions sensitivity (or TPR) and specificity (or 1-FPR), and one using the weighted definitions of sensitivity and specificity (see Equations 12 and 13). These visualizations are similar to Figure 4a. The visualizations are generated automatically; text strings are partially overlapping in some cases.

### Supplementary Data 4 | Cell line information

Tab separated Excel table containing information about the 714 cell lines. Specifically, (from left to right in the table), information about the cell lines (ID, name and description), the binary mutation status of the 60 gene mutation features, and the IC50s for each of the 142 drugs.

## Acknowledgements

TAK would like to thank Bob Duin for fruitful discussions. TAK and FI would like to thank Julio Saez Rodriguez for fruitful discussions. TAK would like to thank Tycho Bismeijer for help with the Python implementation.

